# The structure of an ancient genotype-phenotype map shaped the functional evolution of a protein family

**DOI:** 10.1101/2025.01.28.635160

**Authors:** Santiago Herrera-Álvarez, Jaeda E. J. Patton, Joseph W. Thornton

## Abstract

Some phenotypes are more likely to be produced by mutation than others, but the causal role of these propensities in the evolution of extant phenotypic diversity remains unclear. There are two major challenges: it is difficult to separate the effect of the genotype-phenotype (GP) map from that of natural selection in causing natural patterns of diversity, and most extant phenotypes evolved long ago in species whose GP maps cannot be recovered. Using reconstructed ancestral transcription factors as a model to address this problem, we created libraries containing all possible amino acid combinations at historically variable sites in the proteins’ DNA binding interface (the genotypes) and measured their capacity to bind specifically to response elements containing all possible combinations of nucleotides at historically variable sites in the DNA (the phenotypes). The ancestral proteins we used existed during an ancient phylogenetic interval when a new phenotype—specificity for a new response element—evolved. We found that the two ancestral GP maps were strongly anisotropic (the distribution of phenotypes encoded by genotypes is highly nonuniform) and heterogeneous (the phenotypes accessible around each genotype vary dramatically among genotypes), but the extent and direction of these properties differed between the maps. In both cases, these properties steered evolution toward the lineage-specific phenotypes that evolved during history. Our findings establish that ancient properties of the GP relationship were causal factors in the evolutionary process that produced the present-day patterns of functional conservation and diversity in this protein family.

## MAIN TEXT

Countless conceivable lifeforms have evolved rarely or never, and those that exist are mostly restricted to specific lineages^1–4^. No flying vertebrates have two pairs of wings, for example, and no turtles or frogs fly. What explains the distribution of phenotypes in nature? Classical explanations focus on the influence of selection^5,6^, but the propensities of biological systems to produce phenotypic variation could also shape evolutionary outcomes. A phenotype can become fixed in an evolving population only if it is first generated by mutation. If biological systems are more likely to produce some phenotypes than others^7–13^, and if these propensities change over time as lineages diverge^14,15^, then some phenotypes will be more likely to evolve in some taxa than in others, even in the absence of selection.

Characterizing the impact of phenotype production on evolution is challenging, because most patterns of phenotypic variation observed in nature could arise from production propensities, natural selection, or both, and disentangling their past influences is difficult^12,16–19^. Ideally, we would isolate the phenotype production process by directly characterizing the complete genotype-phenotype (GP) map, which maps all possible combinations of mutations to the phenotypes they encode. This would allow us to precisely quantify the ability of a genetic system to produce phenotypic variation, both on a global scale and by mutation from each particular genotype. But the number of possible genotypes and phenotypes is far too vast, so characterizing GP maps requires reducing its dimensionality while maintaining combinatorial completeness. We reasoned that this goal should be achievable for functionally important sites within a protein and for particular biochemical phenotypes that mediate the protein’s biological functions. The scope of relevant genetic variation can be defined as all combinations of all 20 amino acid states at sequence sites that determine the protein’s phenotype of interest^20–24^. This kind of combinatorial library of protein variants can be engineered and experimentally characterized using deep mutational scanning (DMS)^25^. Combinatorial DMS studies to date have assayed only one or a few phenotypes that exist in extant proteins^24^, so they cannot directly address the role of the GP map in evolution, which requires characterizing the propensity of mutations to produce not only those phenotypes that evolved but also those that did not. The scope of relevant biochemical phenotypes can be delimited as the ability of a protein to carry out its core biochemical activity on all possible substrates of a defined class^26–30^; for a transcription factor, for example, binding could be measured to oligonucleotides containing all combinations of nucleotides at a defined set of sites, such as those that vary among the binding sites for the extant proteins.

The phenotypes of extant lineages evolved long ago, so understanding the causal role of phenotype production in historical evolution requires GP maps to be characterized as they existed in the deep past. Ancestral protein reconstruction^31^ can address this problem by inferring the sequences of ancient proteins using statistical phylogenetic methods; the reconstructed protein can then be used as the background for a DMS experiment. Characterizing GP maps across a phylogenetic time series of reconstructed ancestral proteins^22,32^ would reveal how phenotype production propensities may have changed over time and whether these propensities are congruent with the trajectories of phenotypic evolution that actually unfolded during history.

Here we apply this approach to assess how the structure of the GP map shaped functional diversification in the steroid hormone receptor protein family. We experimentally characterize GP maps of the binding interface of two reconstructed ancestral steroid hormone receptor DNA binding domains (SR DBDs) and their ability to encode specific recognition of all possible DNA response element sequences, with the scope limited to sites in the protein-DNA interface that varied during historical evolution. We then analyze these maps to understand 1) how they could shape potential phenotypic outcomes of evolution on short and long timescales, 2) characterize the mechanisms that changed key features of the maps across evolutionary time, and 3) assess the impact of the maps on the historical evolutionary processes that yielded the lineage-specific patterns of DNA specificity in extant steroid hormone receptors.

### Two ancestral GP maps

SRs are a family of transcription factors that regulate physiological and reproductive biology in bilaterian animals. Most bilaterian taxa have a single SR, which specifically binds to inverted palindromes of the motif AGGTCA, called the estrogen response element (ERE; Fig. 1a). In chordates, a gene duplication of the ancestral SR (AncSR1) produced two major SR classes, which have different DNA specificity phenotypes: chordate estrogen receptors (ERs) retain the ancestral ERE specificity, but a novel specificity for a palindrome of AGAACA, called the steroid response element (SRE), evolved in the lineage leading to AncSR2, the common ancestor of the chordate ketosteroid receptors (kSRs; Fig. 1a, b)^33^. Specificity for DNA is determined primarily by the amino acid sequence of a recognition helix (RH) that binds in the DNA major groove^34,35^. AncSR1 and AncSR2 DBDs differ by 34 amino acid replacements, but experiments on the reconstructed proteins established that three amino acid changes in the RH were the primary cause of the evolution of SRE specificity^33^.

**Fig. 1.**
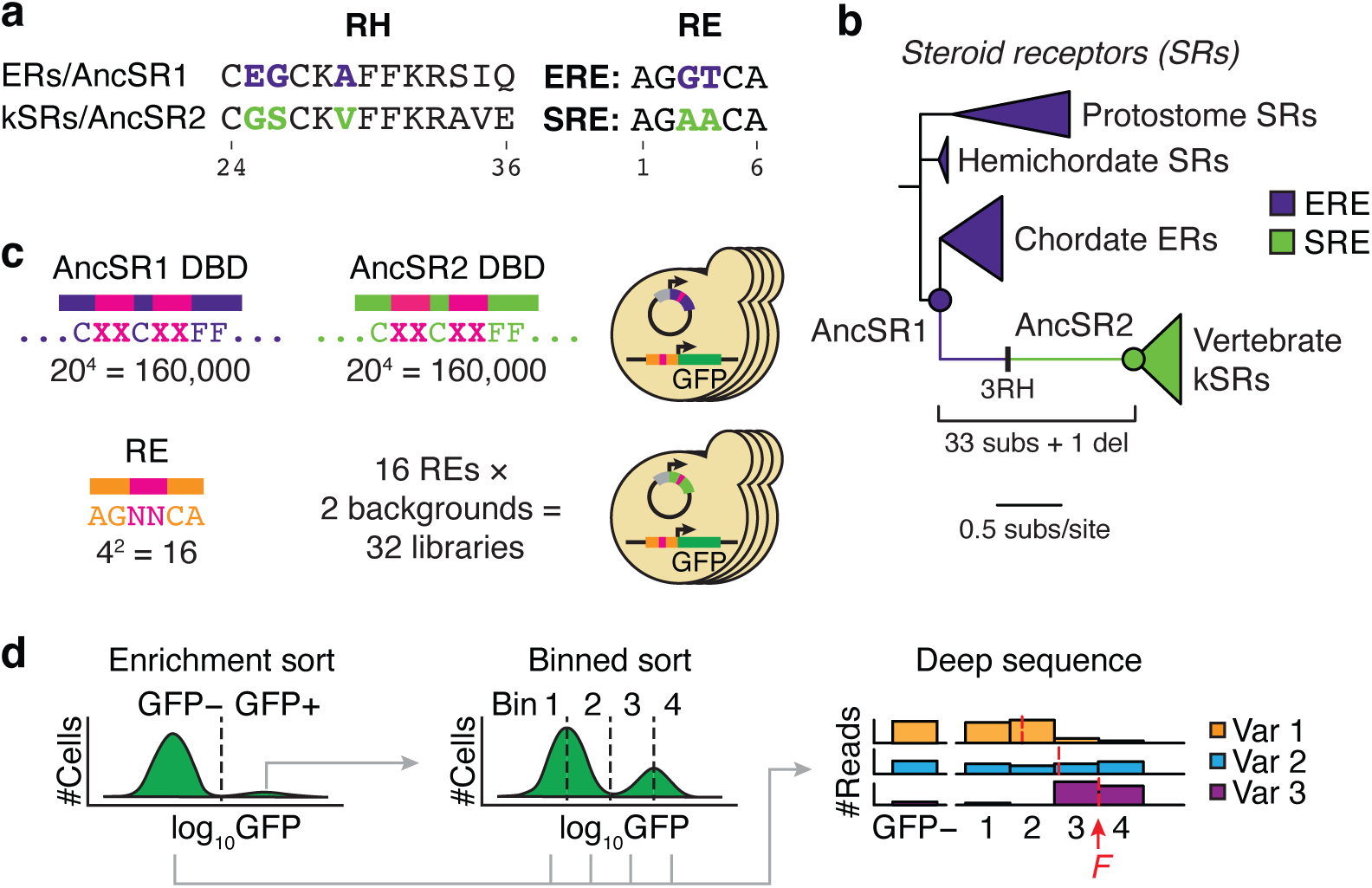
Characterizing ancestral GP maps using multi-phenotype DMS. **a**, Amino acid sequence of the recognition helix (RH) in extant and ancestral steroid receptor (SR) proteins and the sequence of the RE they bind to. Colored residues are responsible for differences in protein-RE specificity. **b**, Phylogeny of SRs. Each clade of proteins is colored by the RE sequence it recognizes. In chordates, a historical transition from ERE to SRE specificity occurred along the branch between AncSR1 (the common ancestor of all chordate SRs) and AncSR2 (the common ancestor of vertebrate kSRs). The number of historical sequence changes along the AncSR1-AncSR2 branch is shown; three of these in the recognition helix (RH) caused the specificity switch^33^. **c**, **d**, DMS experiment to assay effects of RH genotype on binding to variable REs. **c**, We built combinatorial libraries of all combinations of 20 amino acid states at four variable sites in the RH (pink Xs), using the rest of the AncSR1 and AncSR2 DBDs as backgrounds (top left). These were transformed into 16 *S. cerevisiae* strains, each containing one of the 16 possible RE motifs (pink Ns, bottom left) genomically integrated upstream of a GFP reporter gene (right). **d**, We assayed binding of DBD-RE complexes using FACS coupled with deep sequencing. For each library, we performed an initial enrichment sort to select for GFP+ cells. We then grew up the selected cells, pooled them across the 32 libraries, and resorted them into four fluorescence bins in triplicate (binned sort). Sorted cells were deep sequenced to estimate the mean log_10_GFP (*F*) of each combination of protein and RE genotypes.

To understand how phenotype production may have shaped the evolution of SR-DBD specificity, we characterized combinatorially complete GP maps of the DBD-response element (RE) interface at the key ancestral timepoints AncSR1 and AncSR2. The scope of genotypes is all possible 20^4^ = 160,000 amino acid variants at four variable sites in the recognition helix—the three that changed between AncSR1 and AncSR2, plus one other that varies in the broader nuclear receptor family (Fig. 1c). The scope of specificity phenotypes consists of all 4^2^ = 16 possible RE sequences that can be produced by all combinations of nucleotides at the two base positions that vary between ERE and SRE; although extant SRs are not known to bind to many of these REs, there is no a priori biophysical reason that any of them should be impossible to bind. These two maps of the recognition helix-RE interface can be thought of as submaps within the much larger GP map of the entire DBD, which are connected by the 31 other “background” substitutions that occurred between the AncSR1 and AncSR2 proteins (Fig. 1b).

We engineered two protein libraries, each containing all 160,000 variants of the recognition helix in the background of either the AncSR1 or AncSR2 DBD, along with 16 yeast strains, each containing a GFP reporter driven by one of the REs (Fig. 1c, Extended Data Fig. 1a–e). We transformed each RE strain separately with the two protein libraries, with barcodes to mark the strain and the ancestral background, for a total of 5.12 million protein-DNA complexes. We used an initial round of fluorescence-activated cell sorting to enrich the yeast libraries for GFP-positive cells, pooled the enriched libraries, sorted cells in three replicates by their fluorescence, and sequenced the sorted bins (Fig. 1d, Extended Data Fig. 1f, g). Using this strategy, we obtained empirical fluorescence estimates for the majority of complexes with good replicability (*r*^2^ = 0.92 across replicates, excluding complexes at the lower bound of fluorescence; Extended Data Fig. 2). Fluorescence of the remaining complexes was predicted using a generalized linear model trained on the experimental data (Methods, Extended Data Fig. 3a–d)^36,37^.

Each protein variant was assigned a DNA specificity phenotype based on these experiments. A protein variant is classified as specific if it is functional in complex with only one RE, promiscuous if it is functional on multiple REs, or nonfunctional if it is not functional on any RE. We defined functional complexes as those having fluorescence at least as great as the wild-type complex in each background (*i.e.* EGKA:ERE for the AncSR1 library and GSKV:SRE for AncSR2) (Methods, Extended Data Fig. 3e–g).

### Anisotropy in the AncSR1 GP map

The probability that a phenotype will evolve equals the probability that it will be produced by mutation times the probability that, once produced, it will be fixed. The potential impact of the GP map on evolutionary outcomes can be understood by comparison to a null scenario in which the map has two properties: isotropy—all phenotypes are present with equal frequency across the map as a whole—and homogeneity—all starting genotypes in the map produce the same distribution of phenotypes by mutation^38–41^. In that case, the frequency distribution of phenotypes produced by genetic variation would be uniform at both local and global scales, so the outcomes of phenotypic evolution would be affected only by selection and drift. By contrast, anisotropy in the GP map would make some phenotypes more likely to be produced than others, increasing the probability that they will evolve relative to their probability given an isotropic map. Heterogeneity would cause the probability that each phenotype will be produced—and so too the probability that it will evolve—to change as lineages diverge from each other across the map. The extent and direction of anisotropy and heterogeneity define the potential impact of the GP map on evolutionary outcomes.

We characterized the extent and direction of anisotropy in the AncSR1 GP map by measuring the frequency distribution of DNA specificity phenotypes encoded by all functional protein variants. Only 107 out of 160,000 total genotypes in the library were functional (0.07%). Of these, the majority (91 genotypes) were specific for a single RE. We quantified anisotropy using the metric *B,* which expresses the phenotype frequency distribution’s deviation from uniformity as 1 minus the Shannon entropy (base 16); *B* can range from 0 when specificity for all 16 REs is encoded with equal frequency to 1 when only a single phenotype is encoded. We found that the global production distribution is strongly anisotropic (*B* = 0.42). Two specificity phenotypes—ERE and SRE—together account for >60% of all specific genotypes, and only five others can be produced at all; nine phenotypes are not encoded by any protein variant (Fig. 2a).

**Fig. 2.**
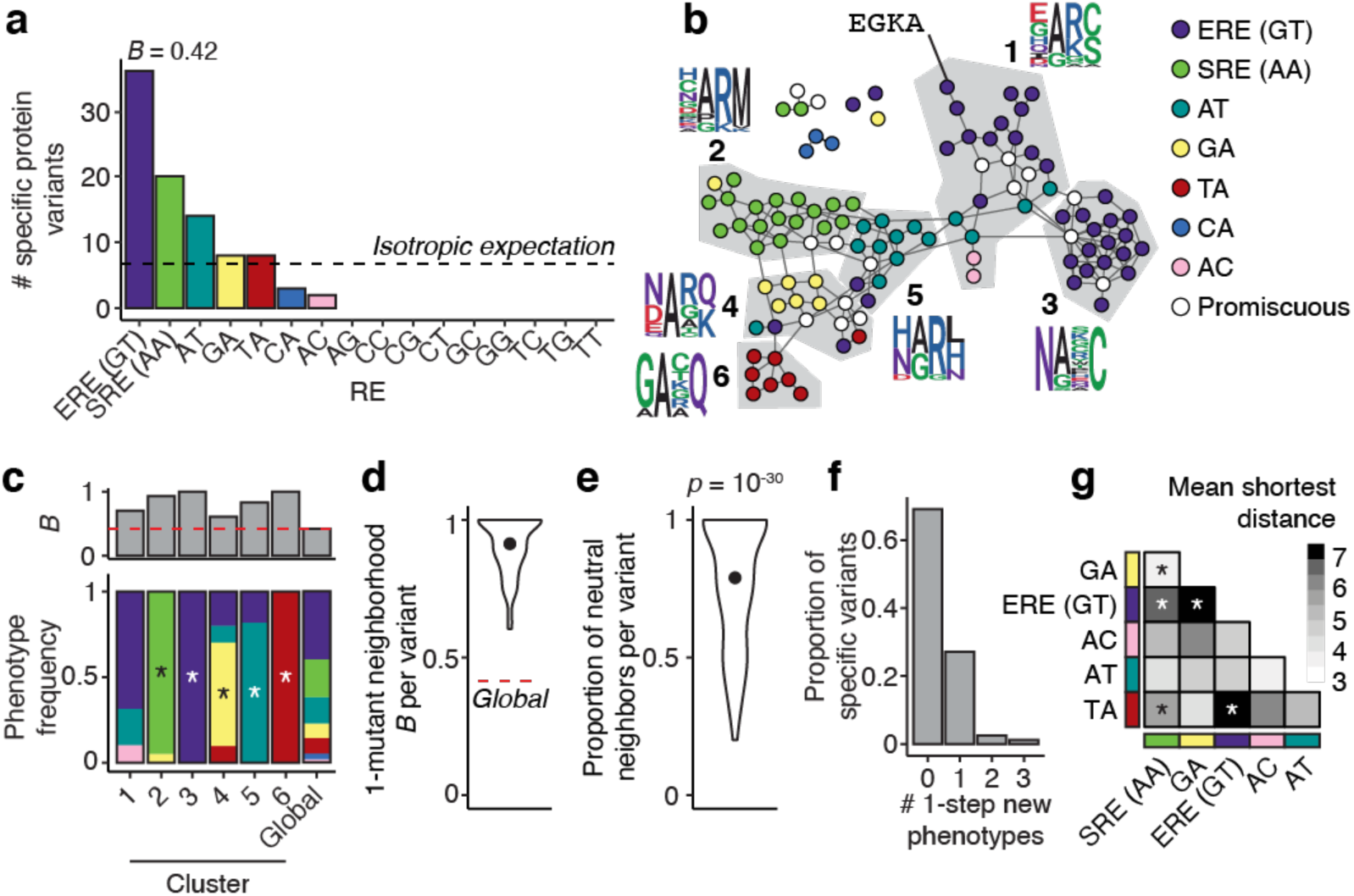
Anisotropy and heterogeneity in the AncSR1 GP map. **a**, Global production distribution of phenotypes in the AncSR1 GP map. Bars represent the number of protein variants that bind specifically to each RE. The dashed line shows the expected frequencies if the distribution were uniform. *B,* the strength of anisotropy, is calculated as one minus the entropy (base 16) of the distribution. **b**, Sequence space network of the AncSR1 GP map. Nodes represent functional protein variants, colored by their RE specificity; white nodes, promiscuous genotypes. Edges connect protein variants that can be interconverted by a single nucleotide change. Genotype clusters (1–6, ordered by decreasing size) identified by a community structure detection algorithm are shown in gray. Sequence logos show amino acid frequencies at the variable RH sites in each cluster. **c**, Bottom: Frequencies of specificity phenotypes within each genotype cluster; the global production distribution is shown for comparison. Asterisks, phenotypes significantly enriched within a cluster relative to the global production distribution (one-sided Fisher’s exact test, *p* < 0.05 after Bonferroni correction). Top: nonuniformity of phenotype production (*B)* within in each cluster. Red line, *B* of global production distribution. **d**, Distribution of *B* of the 1-mutant neighborhood of every RE-specific protein variant in the main network component. Dot shows the mean. Dashed red line, global anisotropy *B* calculated across all genotypes. **e**, Proportion of neutral neighbors per RE-specific protein variant in the main component of the AncSR1 map. Dot shows the mean. *P-value,* probability that the mean would be at least as great as observed if phenotypes were randomly reassigned in the main component (*n* = 91). **f**, Distribution of the number of new phenotypes accessible within one mutation, across all RE-specific variants in the AncSR1 main component. **g**, Mean distance between pairs of phenotypes in the AncSR1 main component. The color of each cell shows the mean of the length of the most direct path from every genotype encoding one phenotype to every genotype encoding the other. Bonferroni corrected *p*-values for a two-sided permutation test where phenotype associations were shuffled within the main component: * *p* < 0.001.

Anisotropy in the AncSR1 map imposes hard limits on phenotypic evolution. The majority of the 16 possible phenotypes could never evolve in this map, even if they conferred strong fitness advantages, because they cannot be produced by any genotype. The direction of anisotropy is also congruent with evolutionary history: the phenotypes that evolved historically in the two lineages descending from AncSR1 are also the most frequently encoded by genotypes in the map.

### Heterogeneity in the AncSR1 GP map

We next assessed the homogeneity of the AncSR1 GP map using Maynard-Smith’s classic network model of sequence space^42^. Each functional protein variant is a node with its experimentally defined phenotype. Nodes are connected by edges if their amino acid sequences can be interconverted by a single nucleotide change given the standard genetic code. Nonfunctional variants are excluded from the network, based on the assumption that they will be removed quickly from evolving populations by purifying selection.

We found that the distribution of phenotypes in AncSR1 sequence space is strongly heterogeneous. Although the majority of functional genotypes (91%) and phenotypes (6 of 7) are mutually connected in a single main network component, each phenotype tends to be sequestered in a local region (Fig. 2b). Using a community structure detection algorithm^43^, we found that the main network component can be partitioned into six clusters of genotypes that have dense connectivity within clusters and weak connectivity between (Fig. 2b). The deviation of the phenotype frequency distribution from uniformity *B* within every cluster is higher than that of the global production distribution, and 5 of 6 clusters are significantly enriched for a single specificity phenotype, which differ among all 5 clusters (Fig. 2c). The clumpy distribution of phenotypes in sequence space arises from the simple fact that similar genotypes, which are connected to each other in sequence space, are likely to encode similar phenotypes (Fig. 2b).

Because of this heterogeneity, the propensity to produce phenotypes depends strongly on the particular genotype occupied at the time. The one-mutant neighborhood around every genotype is extremely nonuniform (mean *B* = 0.91; Fig. 2d), indicating that individual genotypes can access much less phenotypic variation than is encoded across genotype space as a whole. Most mutations are phenotypically neutral (79% of edges; Fig. 2e), and most genotypes can directly access at most one new phenotype (Fig. 2f). The historical starting genotype (EGKA), for example, has access to only one functional neighbor, which also has ERE specificity. Another consequence of heterogeneity is that phenotypes, aggregated over the genotypes that encode them, are differentially accessible to each other, with substantial variation in the number of mutations required to transform each phenotype into the others (Fig. 2g). For example, SRE-specific protein genotypes are directly accessible from nodes encoding specificity for AT and GA, but they are multiple substitutions away from all ERE-specific genotypes, which can reach SRE-specificity only via intermediates with these phenotypes (Fig. 2b).

The GP map of AncSR1 is therefore strongly anisotropic and heterogeneous, and these properties shape the phenotypic variation that can be produced by mutation. Anisotropy across the GP map as a whole favors production of the historical phenotypes ERE and SRE and entirely prevents the production of most conceivable phenotypes. Heterogeneity further restricts the number of accessible phenotypes from each particular genotype, favoring conservation over the evolution of new phenotypes, including conservation of ERE specificity from the historical genotype EGKA.

### The GP map shapes phenotypic outcomes of evolution

To characterize how the structural properties of the AncSR1 GP map could influence the outcomes of evolution, we modeled evolution on the network of functional amino acid genotypes as a discrete-time Markov chain from every possible starting genotype given a variable trajectory length. Each time-step in a trajectory is an amino acid substitution, the probability of which is weighted by the number of single-nucleotide mutations that can mediate it; the relative probability of evolving a given phenotype at the end of the trajectory is the sum of the probabilities of evolving all genotypes that encode it. This model, in which all functional phenotypes have equal fitness, corresponds to neutral molecular evolution with purifying selection^42,44^: it represents a null scenario in which the fixation process does not shape the distribution of phenotypic outcomes except to prevent the loss of function. Using this model allows us to isolate the influence of phenotype production propensities imposed by the GP map’s structure.

We first computed the equilibrium distribution of phenotypic outcomes after an infinite number of substitutions. This represents the limiting case at which the distribution of outcomes is insensitive to the starting genotype and does not change with additional substitutions. The equilibrium outcome distribution is well correlated with the global frequency distribution of phenotypes (Fig. 3a, *r*^2^ = 0.82), reflecting the map’s anisotropy. However, there are differences: the equilibrium distribution is even more nonuniform than the global production distribution (*B* = 0.46), and ERE and AT specificity are the most likely equilibrium outcomes, whereas ERE and SRE specificity are the two most frequently encoded phenotypes (Fig. 3a). This difference arises because most AT-specific genotypes are located centrally within the network, while SRE-specific genotypes are in a more peripheral cluster (Fig. 2b) and are therefore less likely to be occupied. The heterogeneous connectivity of functional genotypes and anisotropy in phenotype production therefore strongly influence evolutionary outcomes, even over infinitely long timescales.

**Fig. 3.**
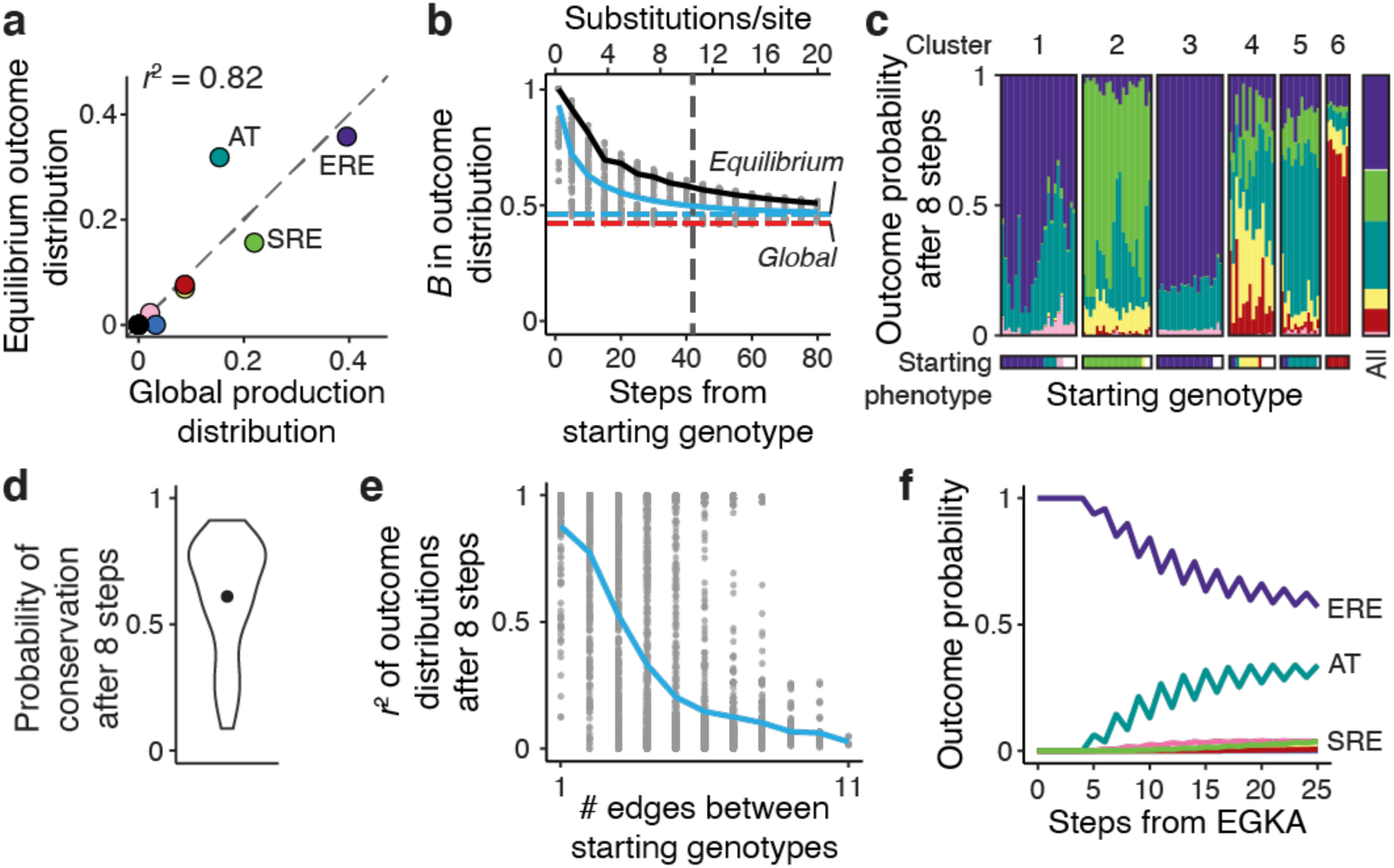
The AncSR1 GP map favors phenotype conservation in evolutionary outcomes. **a**, Comparison between the global production distribution of phenotypes in the main network of the AncSR1 GP map and the long-term equilibrium distribution of phenotypic outcomes of evolutionary trajectories across that map. Each dot shows the frequency of one specificity phenotype in the two distributions. Black dot at the origin represents nine phenotypes not encoded in the map. Dashed gray line, *y* = *x*. Squared Pearson’s correlation coefficient is shown. **b**, Nonuniformity (*B*) in evolutionary outcomes as a function of the length of evolutionary trajectories. Each gray dot shows the *B* of the outcome distribution for trajectories of a given number of substitutions starting from one node on the main network component. Solid blue and black lines show the mean across all starting genotypes and from EGKA, respectively. Dashed horizontal red and cyan lines show *B* of the global production distribution and the equilibrium distribution, respectively. Vertical dashed line shows the number of substitutions required for mean *B* to reach within 0.05 units of the equilibrium value. The secondary *x-*axis (above) shows the trajectory length as substitutions per site. **c**, Distribution of evolutionary outcomes after 8 substitution steps from every starting genotype in the AncSR1 main network component, organized by the cluster of the starting genotype (top). Bottom bar shows the phenotype of each starting genotype. Bars at right show the average outcome distribution for all starting genotypes. **d**, Distribution of the probability of phenotype conservation after 8 substitution steps across all specific starting genotypes in the AncSR1 main network component. Dot shows the mean. **e**, Evolutionary outcomes become less similar as starting genotypes diverge from each other. Each dot shows the similarity of the distributions of phenotypic outcomes (Pearson’s *r*^2^) of 8-step trajectories starting from a pair of genotypes, versus the number of network edges between the pair. Blue line, mean similarity across all pairs of starting genotypes. **f**, Probability of evolving each specificity phenotype starting from EGKA as a function of the number of substitutions.

On finite timescales, heterogeneity strongly affects evolutionary outcomes. After three substitutions, for example—the shortest path between the historical ancestral and derived genotypes—the outcome distributions are highly nonuniform (mean *B* = 0.8 across starting genotypes, Fig. 3b), because most genotypes can reach only a few new specificity phenotypes by a path of this length (Extended Data Fig. 4a). The effect of local heterogeneity on outcomes gradually decays as trajectories get longer, but it takes 42 substitutions (10.5 per site) for the mean *B* to decrease to within 0.05 units of the equilibrium (Fig. 3b, vertical dashed line). By comparison, the maximum root-to-tip branch length in the steroid receptor DBD phylogeny (Fig. 1b), which spans over 500 million years of evolution, is just 2.2 substitutions per site. The phenotypes likely to evolve on phylogenetically relevant timescales are therefore strongly affected by heterogeneity in the GP map.

Another consequence of heterogeneity is that outcomes are strongly contingent on the genetic starting point. Consider a trajectory length of eight substitutions—long enough for new phenotypes to become accessible from most starting points, but not so long that the influence of local heterogeneity is lost. At this timescale, genotypes differ dramatically in the distribution of phenotypes that evolve from them (Fig. 3c). Much of this variation is explained by the genotype cluster to which the starting node belongs (Fig. 3c), because evolutionary trajectories rarely jump between weakly connected clusters and clusters are strongly enriched for individual phenotypes. Even at this timescale, the structure of the GP map favors conservation of the starting phenotype on average (Fig. 3d), but when new phenotypes evolve, those that are acquired also differ strongly among starting genotypes (Fig. 3c).

A final consequence of heterogeneity in the GP map is that as lineages diverge from each other across the map, the distributions of phenotypic outcomes likely to evolve from them become increasingly dissimilar among lineages. The correlation between the distributions of phenotypic outcomes after eight-step evolutionary trajectories from pairs of starting genotypes depends strongly on the distance between those genotypes in the network. For pairs of genotypes that are one substitution apart, the average *r*^2^ is 0.88, but this correlation drops to 0.50 when the genotypes are three steps apart and is entirely lost at 11 steps (*r*^2^ = 0.02, the maximum distance on the network) (Fig. 3e). The likely outcomes of phenotypic evolution therefore become distinct among lineages as they traverse the GP map.

### The AncSR1 GP map favored historical conservation of ERE specificity

Anisotropy and heterogeneity have a very strong and long-lasting impact on the outcomes of evolutionary trajectories that begin from the historical genotype of the recognition helix in AncSR1 (EGKA). It takes 80 substitutions for the nonuniformity in phenotypic outcomes from this starting point to decay to within 0.05 units of equilibrium, almost double the average across genotypes (Fig. 3b, blue vs. black solid lines). It takes at least five substitutions for any new specificity phenotype to be accessed, and even after eight substitutions the probability of conserving ERE specificity is still 0.90 (Fig. 3f). The AncSR1 GP map heavily favors phenotypic conservation from the historical starting genotype across phylogenetically relevant timescales. Unequal phenotype production propensities imposed by the GP map are therefore congruent with the long-term historical conservation of ERE specificity in the lineages that descend from AncSR1 and lead to modern-day estrogen receptors.

The historical outcome that evolved in AncSR1’s other descendant linage—acquisition of SRE specificity in the kSR clade—was very unlikely on phylogenetic timescales. SRE-specific genotypes are distant from EGKA (Fig. 2b), so the probability of evolving SRE specificity after eight substitutions is only 0.0008 (Fig. 3f), despite the fact that this is the second-most frequently encoded specificity phenotype in the network overall. The only specificity phenotype with even moderate probability of evolving at this timescale is AT (probability 0.08; Fig. 3f). Heterogeneity in the genetic neighborhood around EGKA therefore overrides the global propensity for SRE specificity, making the historical outcome that occurred in the kSR clade extremely unlikely.

### Evolution of a different GP map in AncSR2

Given that SRE specificity was unlikely to be produced by mutation from the ancestral genotype in the AncSR1 map, how could this phenotype have historically evolved in the kSRs? We reasoned that the GP map must have changed along the branch leading to AncSR2 on which SRE specificity was acquired. Previous experiments showed that the background substitutions that occurred outside the recognition helix during this interval had a nonspecific permissive effect on both ERE and SRE activation, which allowed the protein to tolerate the historical substitutions and other mutations in the RH (Fig. 1b)^22,33^. We predicted that the background substitutions had a similarly permissive effect across all 16 REs, increasing the number of functional genotypes in the map and the number of phenotypes they encode.

To assess this hypothesis, we characterized the GP map of the RH sites in AncSR2 and compared it to the map in the AncSR1 background. As predicted, the number of functional genotypes and phenotypes both massively increased (Fig. 4a, b). There are 2,407 functional protein genotypes in the AncSR2 map, an increase of >20-fold over the AncSR1 background. 14 of the 16 possible specificity phenotypes are now encoded, twice as many as in the AncSR1 background (Fig. 2a, 4a). The background substitutions therefore dramatically expanded the functional genetic and phenotypic variation that can be produced within the recognition helix.

**Fig. 4.**
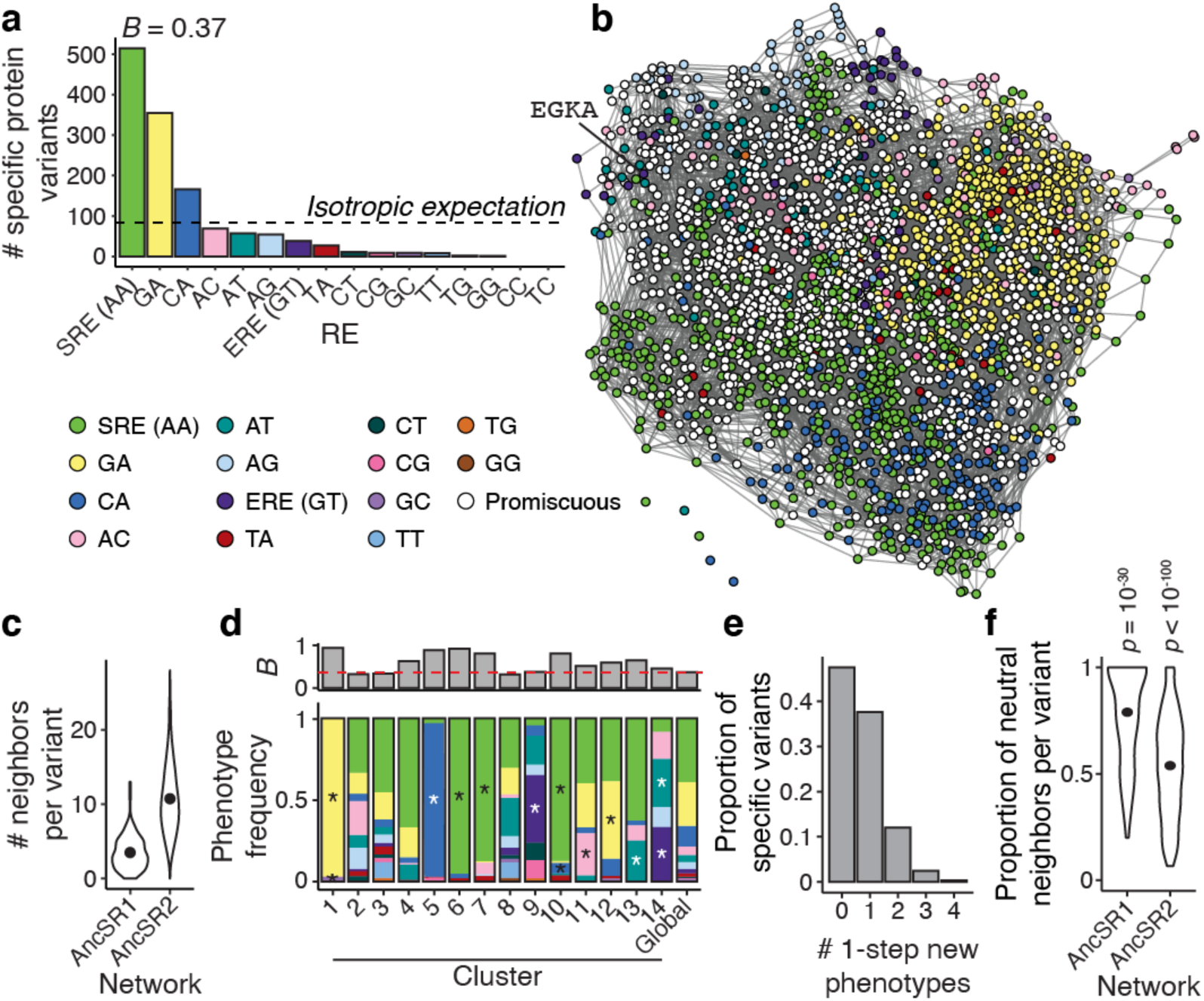
Anisotropy and heterogeneity changed in the AncSR2 GP map. **a**, Global production distribution and global *B* of the AncSR2 GP map. **b**, Sequence space network of the AncSR2 GP map. **c**, Number of one-step neighbors per protein variant in each network. Dots show the mean of each distribution. **d**, Bottom: Frequencies of specificity phenotypes within each genotype cluster (1–14, ordered by decreasing size); the global production distribution is shown for comparison. Only the 14 largest clusters, which contain >90% of genotypes, are shown. Asterisks, phenotypes significantly enriched within a cluster relative to the global production distribution (one-sided Fisher’s exact test, *p* < 0.05 after Bonferroni correction). Top: strength of phenotype uniformity (*B*) in each cluster. Red line, *B* of global production distribution. **e**, Distribution of the number of new phenotypes accessible within one mutation, across all RE-specific protein variants in the AncSR2 main component. **f**, Proportion of neutral neighbors per RE-specific variant in the main network component of the AncSR1 and AncSR2 maps. Dots show the mean. *p-value,* probability that the mean would be at least as great as observed if phenotypes were randomly reassigned in the main component of each map (AncSR1 *n* = 91, AncSR2 *n* = 2,402).

Connectivity between genotypes in the map increased, reducing the impact of heterogeneity and facilitating access to new phenotypes. In the AncSR2 network, all but five of the 2,407 functional nodes are connected in a single main component (Fig. 4b), and the mean number of edges per node is 10.7, a three-fold increase compared to the AncSR1 network (Fig. 4c). Genotype clusters are still present, but *B* within clusters is weaker than in the AncSR1 map (Fig. 2c, 4d). As a consequence, genotypes have more access to new phenotypes: >50% of genotypes in the AncSR2 map can access between 1 and 4 new phenotypes within a single mutation (Fig. 4e, compare to Fig. 2e), because genotypes are typically connected to far more non-neutral neighbors (Fig. 4f).

Finally, the global production distribution of phenotypes also changed across this interval. In the AncSR2 map, SRE became the most frequently encoded phenotype (39% of specific variants), and ERE’s rank declined from first to seventh (encoding just 3% of specific variants) (Fig. 2a, 4a). The background substitutions therefore changed the direction of anisotropy in the GP map from favoring the ancestral specificity to producing the derived specificity.

### The AncSR2 GP map favored evolution of SRE specificity

These changes in the AncSR2 GP map dramatically altered the likely phenotypic outcomes of evolution. At long-term equilibrium using our Markov model and the AncGR2 map, the most likely evolutionary outcome is now SRE specificity, with a probability close to 40% (Fig. 5a, compared to <20% in the AncSR1 map). At moderate timescales as well, SRE specificity is the most likely outcome across the majority of starting genotypes (Fig. 5b). The probability of evolving new phenotypes overall is considerably higher in the AncSR2 network compared to AncSR1 (mean probability of conservation after eight steps 0.47 in AncSR2 but 0.61 in AncSR1, Fig. 3d, 5c).

**Fig. 5.**
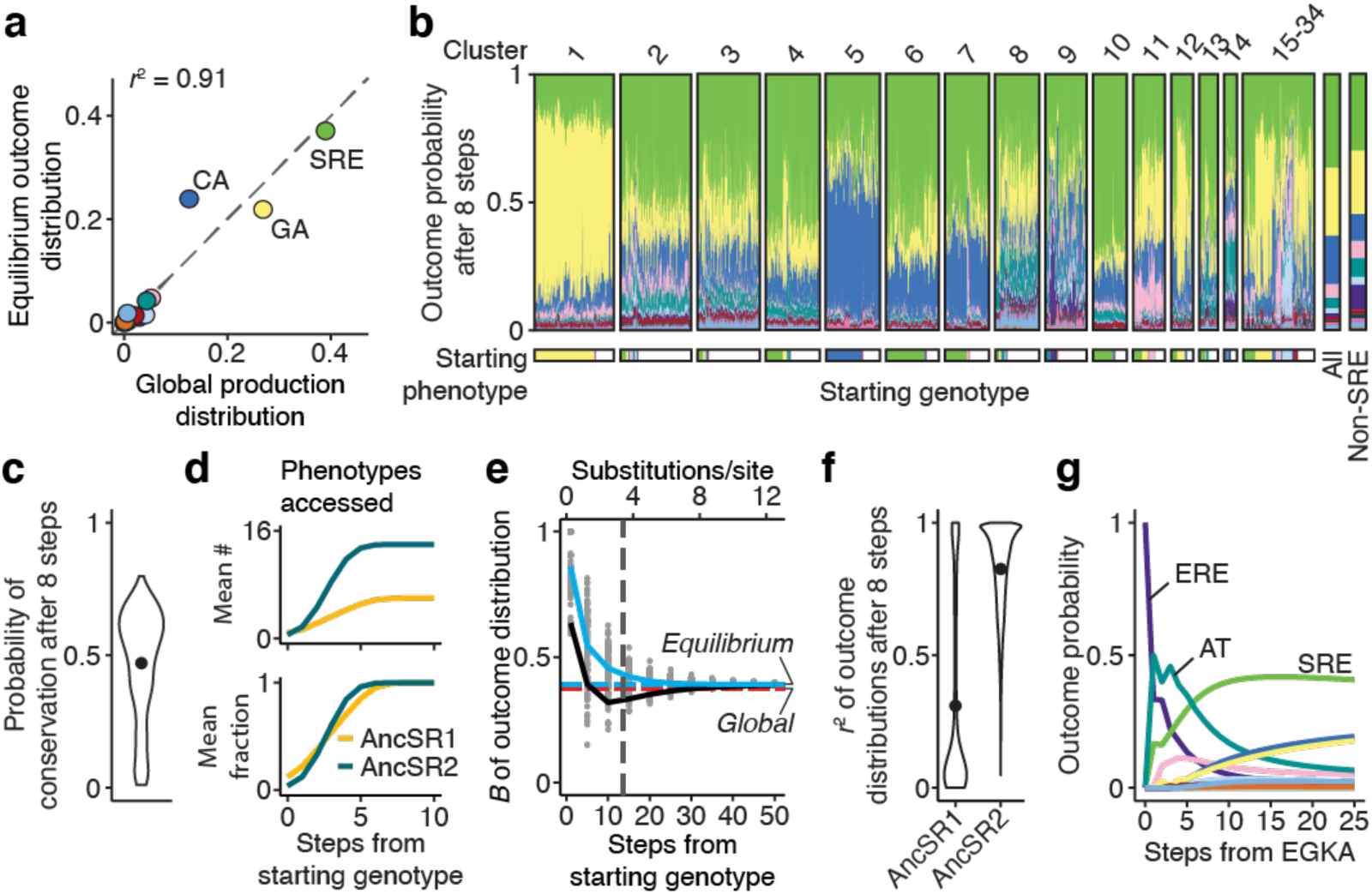
The AncSR2 GP favors the acquisition of SRE specificity in evolutionary outcomes. **a**, Comparison between the global production distribution and the long-term equilibrium distribution of phenotypic outcomes in the AncSR2 main network. Dashed gray line, *y* = *x*. **b**, Distribution of evolutionary outcomes after 8 substitution steps from every starting genotype in the AncSR2 main network component, organized by the cluster of the starting genotype (top). Bottom bar shows the phenotype of each starting genotype. Bars at right show the average outcome distribution for all starting genotypes and all non-SRE-specific starting genotypes, respectively. **c**, Distribution of the probability of phenotype conservation after 8 substitution steps across all specific starting genotypes in the AncSR2 main network component. Dot shows the mean. **d**, Number (top) and fraction (bottom) of phenotypes in each network accessible after evolutionary trajectories of length shown on the *x* axis. Lines show the mean across all starting genotypes in each network. Gold, AncSR1 network; teal, AncSR2 network. **e**, Nonuniformity (*B*) in evolutionary outcomes as a function of the length of evolutionary trajectories. Lines and colors are the same as in Fig. 3b. **f**, Distribution of the similarity in outcome distributions (Pearson’s *r*^2^) for 8-step trajectories starting from all pairs of genotypes in the AncSR1 and AncSR2 main networks. Dots show means. **g**, Probability of evolving each specificity phenotype starting from EGKA as a function of the number of substitutions.

These changes in evolutionary outcomes are attributable to the increased connectivity of the AncSR2 network and the shift in the global production distribution. From every starting point, the increase in functional nodes and connectivity in the AncSR2 network allows evolutionary trajectories to arrive at more genotypes and more phenotypes, and to do so by a smaller number of steps (Fig. 5d, Extended Data Fig. 4b). As a result, the influence of local heterogeneity is lost faster, and trajectories more rapidly converge on the equilibrium distribution (Fig. 5e), which more closely resembles the production distribution than in the AncSR1 background (Fig. 5a). Evolutionary outcomes are also more similar across pairs of starting points than they were in the AncSR1 map (Fig. 5f). Combined with the shift in the global production distribution, this causes SRE specificity—which was already the second-most likely outcome in the AncSR1 map—to become the most likely outcome from a majority of starting points in the AncSR2 background.

From the historical RH genotype EGKA (the AncSR2 protein with the RH states reverted to their ancestral states), the likely outcomes of phenotypic evolution are dramatically different than in the AncSR1 map. EGKA is much less mutationally isolated in the AncSR2 network, so the probability of conserving ERE specificity after eight substitutions drops from 0.9 in the AncSR1 map (Fig. 3f) to 0.07 in the AncSR2 map (Fig. 5g). The probability of evolving new specificity phenotypes on moderate timescales increases accordingly: after just three steps, two new phenotypes—including SRE specificity—are more likely than conservation of ERE. By six steps, SRE specificity becomes the most likely of all phenotypic outcomes. Heterogeneity around the ancestral genotype is likely to have influenced the evolutionary trajectory taken between ERE and SRE specificity, because the vast majority of paths connecting protein variants with these phenotypes pass through intermediates that are AT-specific (Fig. 5g).

The background substitutions that occurred along the branch to AncSR2 therefore changed the GP map of the RH in a way that dramatically changed the probable phenotypic outcomes of evolution. The structural features of this map strongly favor phenotypic diversification and make the particular phenotype that historically evolved in the kSR lineage the most likely of all possible outcomes.

### Simple biophysical mechanisms changed the GP map

Finally, we sought insight into the biophysical mechanisms that changed the structure of GP map of the recognition helix between AncSR1 and AncSR2. Although our experiments provide a functional rather than biophysical readout, different biophysical mechanisms predict different patterns of functional change between the AncSR1 and AncSR2 maps. We therefore analyzed the change in fluorescence of each protein-DNA complex variant between the two backgrounds to identify potential biophysical mechanisms and considered them in light of existing crystal structures. We found evidence for two major mechanisms.

First, the background substitutions between AncSR1 and AncSR2 appear to have caused a universal increase in affinity across all the protein-DNA complexes in our libraries. Previous experiments and crystal structures showed that 11 of the background substitutions improve affinity on both ERE and SRE by generating favorable interactions not involving the two variable RE bases^22,33^. We therefore hypothesized that affinity increased universally for all amino acid variants in the DBD’s recognition helix across all 16 REs. To test this hypothesis, we fit a simple model in which fluorescence in each ancestral background is a function of a complex’s affinity, and affinity is scaled by a constant factor in AncSR2 relative to AncSR1 (Methods). The model fits the data very well (*r*^2^ = 0.95; Fig. 6a), with an estimated 70-fold universal improvement in affinity in the AncSR2 background. This apparent increase in affinity explains the vast increase in functional genotypes and specificity phenotypes between AncSR1 and AncSR2, because many protein-DNA complexes that had weak affinity in the AncSR1 background—and were therefore nonfunctional—bind strongly enough in AncSR2 to produce functional levels of fluorescence. The number of promiscuous protein variants also increases, because many variants cross the threshold for functionality on multiple REs (Extended Data Fig. 5a). A universal improvement in affinity of recognition helix variants across all 16 REs explains not only the increased size but also the greater connectivity of the AncSR2 network: the background substitutions do not qualitatively change the amino acid determinants of binding but instead make them less stringent (Fig. 6b), so many of the newly functional nodes in AncSR2 are close neighbors of those that were already functional in AncSR1, with an average gain of 11 new neighbors per node (Fig. 6c).

**Fig. 6.**
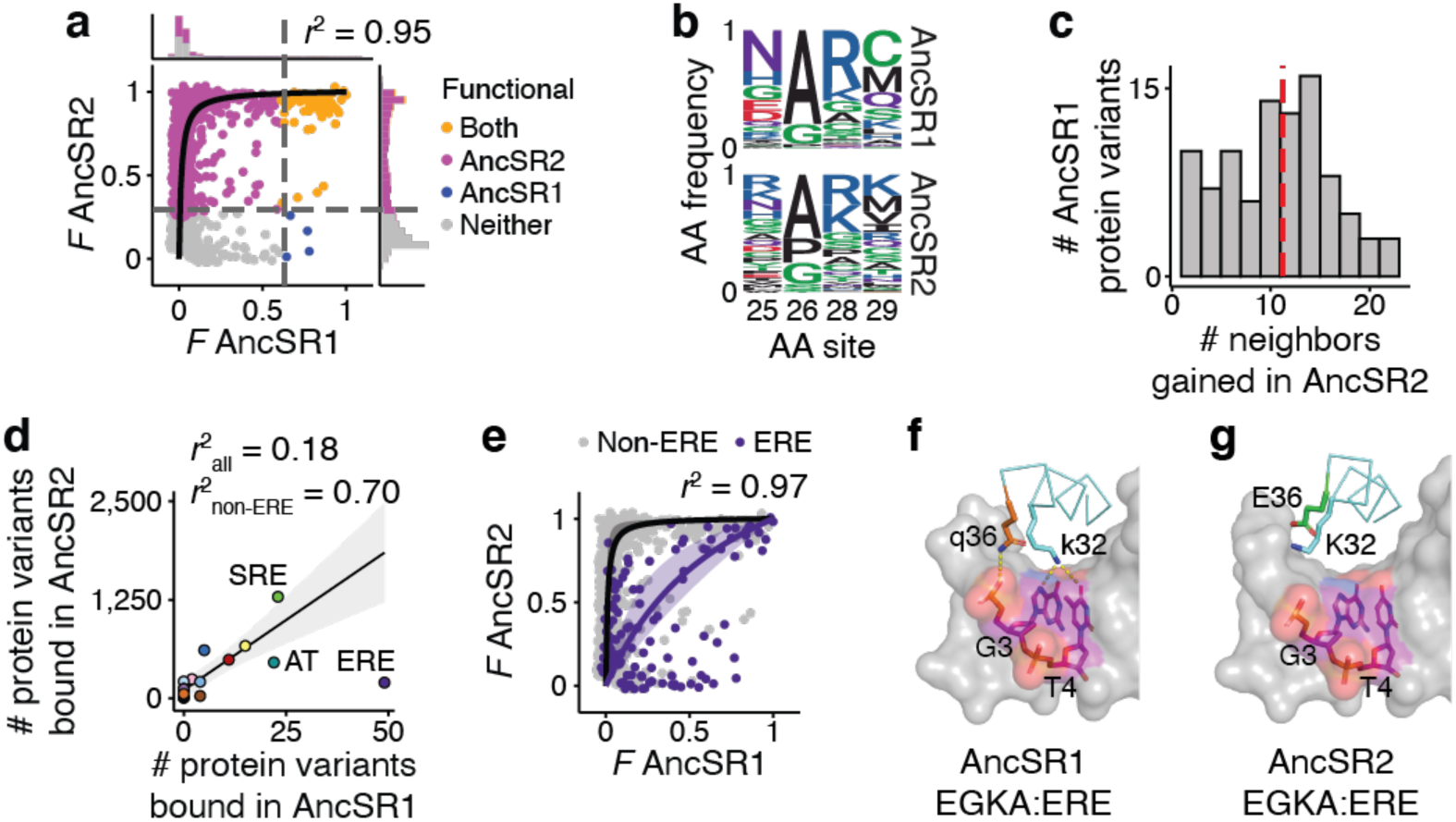
Nonspecific effects of background substitutions on DBD-RE affinity. **a**, Fluorescence of each complex in the AncSR1 vs. AncSR2 background, scaled between the upper and lower bounds for each background. Curve shows best-fit model assuming that the affinity of every complex in the AncSR2 background is related to its affinity in the AncSR1 background by the same scaling factor. Shaded region around the curve (barely visible) shows bootstrapped 95% confidence interval (CI). The Pearson’s *r*^2^ between the data and model predictions is shown (*n* = 2,627). Histograms show distribution of *F* in each background. Dashed lines show the fluorescence of the wild type complex in each background (AncSR1-EGKA:ERE or AncSR2-GSKV:SRE). Colors indicate the backgrounds in which each genotype is functional. **b**, Amino acid frequencies at the variable RH sites across all functional protein variants in the AncSR1 and AncSR2 maps. **c**, Distribution of the number of neighbors gained in the AncSR2 background across all functional protein variants in the AncSR1 background that remain functional in the AncSR2 background. Dashed line, mean. **d**, Correlation between the number of protein variants bound per RE in each background. Black line, linear fit to all REs except ERE; shaded region, 95% CI. **e**, Same as **a**, but fitting a model in which the background substitutions affect affinity of all variants for ERE by one scaling factor and for all other REs by a different scaling factor. Purple, observed fluorescence and best-fit model predictions for ERE complexes; gray, for non-ERE complexes. **f**, Crystal structure of the AncSR1-EGKA protein in complex with ERE (PDB 4OLN). The RH backbone is shown as a ribbon, with key side chains shown as sticks. The gray surface shows ERE, with variable bases and backbone as sticks. In this complex, glutamine (q) at site 36 forms a hydrogen bond (yellow dashed line) with the DNA backbone, and lysine (k) at site 32 forms two hydrogen bonds to the ERE-specific bases G and T. **g**, Same as **f**, but with the AncSR2-EGKA crystal structure (PDB 4OND). Substitution to glutamic acid (E) at site 36 abolishes the ancestral hydrogen bond to the DNA backbone and results instead in electrostatic repulsion from the backbone. This deforms the recognition helix, abolishing the hydrogen bonds between K32 and the G and T bases. In **F** and **G**, lowercase letters represent ancestral amino acid states, and uppercase derived.

The second apparent mechanism is that the background substitutions negatively affect specific binding to ERE, shifting the anisotropy in the global production distribution away from ERE and leaving SRE as the most-encoded phenotype in the AncSR2 background. A universal affinity increase predicts that the number of variants with every specificity phenotype should increase proportionally across the AncSR1-AncSR2 interval; this pattern holds, but ERE is an outlier, with far fewer variants than would be expected given the pattern for other phenotypes (Fig. 6d). Moreover, ERE complexes exhibit notably lower fluorescence in the AncSR2 background than predicted by a universal increase in affinity (Extended Data Fig. 5b). We estimated the effect of the background substitutions on ERE affinity by incorporating a background-by-ERE interaction term into our affinity-fluorescence model; adding this parameter improves the fit to the data (*r*^2^ = 0.97), with the background substitutions improving ERE affinity by an estimated 2.3-fold, compared to 99-fold for all other REs (Fig. 6e). The extent of the relative reduction in fluorescence differs among protein variants, however, suggesting additional specific interactions between background substitutions and amino acids in the recognition helix (Extended Data Fig. 5c). Crystal structures of the EGKA:ERE complex^33^ suggest one possible structural mechanism for the global reduction in ERE affinity: one of the background substitutions (q36E) deforms the protein backbone of the recognition helix, abolishing two hydrogen bonds that are formed between a conserved residue and bases in the ERE (Fig. 6f, g). Corroborating this mechanism, the background substitutions also shift the global production distribution away from AT specificity (Fig. 6d), and this is the only other RE that can form these hydrogen bonds.

The structure of the GP map therefore changed between AncSR1 and AncSR2 via two simple biophysical mechanisms. By increasing the affinity of all protein variants tested for all 16 REs, while also impairing their affinity for ERE, the background substitutions reduced local heterogeneity and changed the direction of anisotropy, facilitating the evolution of many new genotypes and phenotypes and shifting the protein’s global propensity away from conserving ERE specificity to evolving the new specificity for SRE.

### Robustness to assumptions

To assess whether our conclusions are sensitive to assumptions that we made in our analysis, we reanalyzed our experimental data under different models and assumptions. First, we applied different thresholds to classify genotypes as functional or nonfunctional, included promiscuous genotypes when characterizing global production distributions, and characterized these distributions using only genotypes with experimentally measured phenotypes. In every case, we observed similar forms of anisotropy and heterogeneity in both the AncSR1 and AncSR2 GP maps to those reported above (Extended Data Fig. 6).

Second, instead of treating the protein as an evolutionary unit independent of the RE, we repeated our analyses using an alternative sequence space network in which the protein and RE coevolve as a complex. In this model, evolution may occur via single-step amino acid mutations in the protein or nucleotide mutations in the RE. Our main conclusions again hold: anisotropy and heterogeneity impact the outcomes of evolution over long and short timescales, favoring ERE conservation in the AncSR1 map and evolution of SRE specificity in AncSR2 (Extended Data Fig. 7).

Finally, we addressed uncertainty about the ancestral sequences. AncSR1 and AncSR2 DBD reconstructions have very high confidence, containing just five and zero ambiguously reconstructed sites, respectively^32^. Experimental data from a prior single-mutant DMS study show that the effects of mutations in the RH are virtually identical when they are introduced into the AncSR1 background or into an alternative reconstruction of AncSR1 that incorporates all plausible alternative amino acids at the ambiguously reconstructed sites (*r*^2^ > 0.99; Extended Data Fig. 8)^32^. The very limited uncertainty about the AncSR1 ancestral sequence is therefore likely to have little or no effect on our conclusions.

## DISCUSSION

### The GP map was a cause of historical phenotypic evolution

Our data establish that anisotropy and heterogeneity in the two ancestral GP maps studied were causal factors in the historical lineage-specific evolution of DNA specificity. Establishing causality in a multifactorial framework requires 1) evidence that a putative cause increases the probability of the outcome(s) of interest, and 2) evidence for a specific mechanism by which the cause affects the outcome’s probability^45^. Concerning the first requirement, we showed that the particular forms of anisotropy and heterogeneity in the AncSR1 map increased the probability that ERE specificity would be evolutionarily conserved relative to a map that did not have those properties, and those in the AncSR2 map increased the probability that SRE specificity would be acquired. The second requirement is satisfied by a simple axiom of population genetics: the probability that a phenotype will evolve is the product of its probability of production and its probability of fixation under the influence of selection and drift. If the structural properties of the GP map increase the probability that a phenotype is produced relative to a map that does not have these properties, then the probability that the phenotype will evolve must also increase.

A cause must precede its effect. This is unambiguously the case for the anisotropy and heterogeneity that favored conservation of ERE specificity in AncSR1, which was ancestral to the ER lineage in which conservation historically occurred (Fig. 1b). Moreover, the production propensity that favored ERE conservation did not change across hundreds of millions of years of subsequent evolution during which ERE specificity was conserved, because there were zero amino acid changes anywhere in the DBD along any of the descendant branches leading from AncSR1 to ERα in the ancestor of all bony vertebrates. Even most present-day ERα DBDs contain zero or at most a single substitution relative to AncSR1 (Extended Data Fig. 9). As for the acquisition of SRE specificity in the AncSR2 lineage, SRE specificity was already favored in the AncSR1 map as the second-most encoded phenotype. Further, the massive increase in connectivity of the AncSR2 map, which dramatically increased the propensity for new phenotypes to evolve, must have been acquired before SRE specificity actually evolved, because the recognition helix substitutions that conferred SRE specificity during history cannot be tolerated unless the background substitutions that nonspecifically increased DNA affinity occurred first^33^. Our experiments do not resolve whether the third major property of the AncSR2 map—a shift in the direction of anisotropy away from ERE specificity that further enhanced the propensity to encode SRE specificity—occurred before or after this phenotype was historically acquired.

We do not argue that selection played no historical role in the evolution of specificity. It seems likely that purifying selection would have favored conservation of ERE specificity in the chordate ERs, and positive selection could have contributed to fixation of SRE specificity in the AncSR2 lineage. If so, however, selection would have further increased the probability of outcomes that were already favored by the phenotype production propensities caused by the GP map’s structure. Our argument is agnostic to the evolutionary causes of the nonspecific increase in DNA affinity that caused the expansion of the AncSR2: the substitutions outside the RH could have been driven by selection for affinity (not specificity), but they also could have been fixed through a process of systems drift under purifying selection to maintain occupancy on DNA, such as might occur if receptor expression and affinity drifted in opposite directions.

Our analyses show that anisotropy and heterogeneity in the AncSR1 and AncSR2 GP maps can have very strong impacts on evolutionary outcomes. Some phenotype production propensities that we observed are absolute. There are nine specificity phenotypes that cannot be encoded by any genotypes in the AncSR1 GP map, and two cannot be encoded in AncSR2. These phenotypes could never evolve, no matter how large a fitness benefit they might in principle confer, because they cannot be produced. Heterogeneity’s local impacts are also absolute with respect to many phenotypes: from every starting point, the vast majority of phenotypes are impossible to produce directly by mutation, and most require many substitutions before they become accessible. Selection would be powerless to fix these phenotypes over short or medium timescales. The structure of the GP map limited evolution to a small subset of possible phenotypes; history played out within this set.

### Generality

Our experiments addressed just four amino acid sites in the protein-DNA interface, but it seems likely that that the production propensities we observed in our GP maps of the protein’s recognition helix would largely persist in larger maps in which more sites are allowed to vary. DBD crystal structures show that the vast majority of other amino acid sites do not make specific contacts with the variable RE bases^33,35^ and are therefore less likely than the sites we studied to strongly influence the specificity of binding. Incorporating variation at other amino sites could increase or reduce connectivity in the network, altering the effect of heterogeneity within the map on the length of evolutionary trajectories to new phenotypes.

There is evidence that features similar to those we observed in the steroid receptor GP map affect many biological systems and their evolution across levels of organization. Anisotropy is apparent in other molecular^46,47^ and developmental systems^48–51^, and the phenotypes favored by these propensities are often congruent with natural patterns of diversity^52–55^. Heterogeneity also appears to be widespread, because most random mutations are phenotypically neutral if they are tolerated^51,56–59^, and long-term phenotype conservation is widespread in the fossil record^60^. When new phenotypes are acquired, identical perturbations often yield different phenotypes in different lineages^61–64^, and convergent evolution becomes less likely among distantly related lineages^65^. It therefore seems likely that anisotropy and heterogeneity are near-universal characteristics of GP maps^8,39,41^, and that these properties have shaped large-scale patterns of phenotype conservation and lineage-specific evolutionary change across the tree of life.

Life is astonishing in its diversity, but an even deeper puzzle lies in the fact that only a tiny fraction of conceivable phenotypes have ever evolved, and those that have evolved are mostly limited to particular taxa^2–4,66^. Chance and selection are likely to have been important causes of the patchy distribution of phenotypes on Earth, but not the only ones. Our work shows that as a protein or other biological system moves through sequence space, the set of phenotypes that it can produce by subsequent mutations changes at every step. The particular biology that we observe today must therefore reflect the constantly changing potential of diverging biological systems in the past to generate new forms of life at every moment.

### Methods

#### RE reporter strains

To measure binding of SR DBD to the 16 RE variants, we adapted a yeast GFP reporter system previously developed to measure binding to ERE and SRE, where GFP expression is well correlated with DNA affinity over a range of at least 2 M^−2^ (*r*^2^ = 0.74)^32^. We engineered 16 yeast strains, each of which reports on binding of the DBD to one RE. We modified the yeast strain CM997 (YPS1000 MATa ho::KMX)^67^ to replace the *KMX* gene at the *HO* locus with a construct containing yeast-enhanced GFP downstream of a minimal *CYC1* promoter with an array of four palindromic RE sites (tcaAG<u>NN</u>CAcagTG<u>NN</u>CTtga), each separated by a 19-nt sequence, along with a *HygR* gene. To ensure a consistent dynamic range of fluorescence across strains, we made changes to two RE strains in the nucleotide sequences flanking the palindromes at sites that do not affect specificity^34,35^ (see Supplementary Methods for details). These constructs were transformed into yeast using the lithium acetate method^68^ and selected for resistance to hygromycin and susceptibility to G418; integration was confirmed by Sanger sequencing.

To validate this reporter system, we measured fluorescence of each strain in the presence and absence of a DBD variant with universally high affinity to all REs (AncSR1+11P+GGKA)^28,33^. We used a low-copy yeast vector (pDBD) to express this DBD variant as a C-terminal fusion with an SV40 nuclear localization signal and a *S. cerevisiae* Gal4 activation domain (Gal4AD) under control of a pGAL1 promoter. We transformed this construct into each yeast strain using the lithium acetate method followed by G418 selection (50 μg/mL). Single colonies were inoculated in YPD+G418 and transferred to YPGal+G418 media for 6 hours to induce DBD expression. GFP fluorescence was measured on a BD LSRFortessa flow cytometer using a 488 nm laser with 505 nm long pass and 525/50 nm band pass filter. We used as the metric of fluorescence log_10_(GFP/FSC-A^1.5^), which normalizes fluorescence to cell volume. All 16 strains showed DBD-dependent fluorescence across a similar dynamic range (Extended Data Fig. 1a–c).

#### AncSR1 and AncSR2 combinatorial library construction

We used as the wild-type protein sequences the maximum *a posteriori* AncSR1 and AncSR2 DBD sequences inferred from a maximum likelihood phylogeny of nuclear receptors^32^. We optimized codon usage for yeast and cloned the ancestral DBDs into the pDBD2.1 expression vector, which is modified from the pDBD vector^22,32^ to express GFP at a level within the dynamic range of fluorescence for the wild type AncSR1:ERE and AncSR2:SRE complexes. A bidirectional pGAL1/GAL10 promoter simultaneously drives DBD and mCherry expression, which allowed us to monitor plasmid retention in yeast (Extended Data Fig. 1d).

Combinatorial mutant libraries were created by synthesizing oligos (IDT) with degenerate NNS codons to encode all 20 amino acids and a stop codon at four recognition helix sites of each ancestral protein (Extended Data Fig. 1e). To distinguish sequencing reads coming from different RE strains, 16 synonymously barcoded versions of the library were designed for each background (Extended Data Fig. 1e, Supplementary Table 1). Each barcode (REBC) differed by at least three nucleotides to ensure accurate read assignment despite sequencing errors. The oligos were cloned into the pDBD2.1 vector using the BsaI-HF Golden Gate Assembly kit (NEB), transformed into Invitrogen ElectroMAX DH5ɑ-E E. coli, and maxiprepped (Supplementary Methods). Transformation yields exceeded 1.08×10⁷ cfu per barcoded library, providing 56-fold coverage of the amino acid library size (Supplementary Table 2). Assemblies were validated by Sanger sequencing of independent transformants and PCR of the plasmid libraries to confirm the correct insert size.

Maxiprepped libraries (GenElute HP, Sigma-Aldrich) were transformed into the yeast reporter strains using an optimized yeast electroporation protocol (Supplementary Methods). Transformation yields exceeded 10^7^ cfu per library (50-fold coverage), estimated by dilution plating (Supplementary Table 2). Yeast libraries were flash-frozen in liquid N_2_ in 200 OD_600_-mL aliquots with 25% glycerol and stored at –80°C. Multiple transformant rates estimated from Sanger sequencing of individual colonies^69^ were estimated to result in 0.03% or fewer cells with multiple plasmid copies at time of sorting.

#### Cell sorting

We used fluorescence-activated cell sorting (FACS) to separate cells based on their GFP expression. We performed two rounds of sorting: an initial “enrichment sort” to enrich for GFP+ variants in the full libraries, and a second, higher resolution “binned sort” on the enriched libraries to generate quantitative fluorescence estimates for each variant. Enrichment sorting was performed in batches of 8 libraries. Two glycerol stocks per library were thawed on ice, after which cells were recovered for 2 hours in 400 mL YPD+chloramphenicol (chlor) per library at 30°C and 225 rpm. After recovery, G418 was added to the culture and a sample of cells was taken for dilution plating. We recovered a minimum of 1.6×10^7^ cfu per library (82-fold coverage). After 15 hours of overnight growth, libraries were washed once in PBS, resuspended to OD_600_ 0.25 in 50 mL YPGal+G418, and grown for 6 hours to induce DBD expression. Cells were then spun down, washed once in PBS, resuspended in 5 mL PBS, and kept on ice for sorting.

Sorting was performed at the University of Chicago Cytometry and Antibody Technology Facility on a BD FACSAria Fusion machine. We used a 488 nm laser with 495 nm long pass filter and 515/20 nm band pass filter for GFP detection, and a 561 nm laser with 595 nm long pass filter and 610/20 nm band pass filter for mCherry detection. After gating on homogeneous single cells and mCherry expression, we sorted cells into GFP– and GFP+ populations (Extended Data Fig. 1f). To normalize fluorescence to cell volume, GFP gates were drawn to have a slope of 1.5 on a log(FSC-A)-log(GFP) plot. We sorted 2.5×10^7^ cells per library in the enrichment stage (129-fold coverage, Supplementary Table 2).

Enriched cells from different libraries were pooled by GFP bin and grown in either 700 mL (GFP+) or 2 L (GFP–) of YPD+G418+chlor. Cultures were grown overnight at 225 rpm and 22–30°C, depending on the ratio of cells to media, until they were at least OD_600_ 3 but not yet saturated. 200 OD_600_-mL 25% glycerol stocks were then made for both the GFP+ and GFP– cultures. 10 OD_600_-mL of the GFP– culture was used for plasmid extraction using a previously described protocol^25^.

The binned sort was performed to yield three replicates per library. For each replicate, two 200 OD_600_-mL glycerol stocks of GFP+ cells per enrichment sort batch were thawed on ice, recovered in 400 mL YPD+chlor for 2 hours, and sampled for dilution plating. After adding G418, cultures were grown overnight, achieving a recovery rate at least 4X the number of GFP+ cells collected during the enrichment sort (Supplementary Table 3). Overnight cultures were pooled proportionally to the GFP+ cell counts from the enrichment sort, yielding a total of 100 OD_600_-mL. The pooled cells were washed with PBS, induced for DBD expression in 400 mL YPGal+G418 for 6 hours, washed again, resuspended in 40 mL PBS, and kept on ice for sorting. Binned sorting followed the enrichment sort protocol but used four GFP bins instead of two (Extended Data Fig. 1g), with ∼1.6×10⁸ cells collected per replicate. The number of sorted cells and recovered reads was consistent across libraries and replicates (Supplementary Table 4).

#### Deep sequencing

After sorting, cells were grown in 100 mL YPD+G418+chlor per 10⁷ sorted cells, or at least 100 mL per bin. Cultures were grown overnight to at least OD_600_ 3.0 but not yet saturated, and 50 OD_600_-mL was collected per 10⁷ sorted cells for plasmid extraction.

Sequencing libraries were constructed from plasmids extracted from the enrichment sort GFP– population and the four binned sort populations using two rounds of amplification. In the first round, the RH scanning and REBC regions of the DBD were amplified with primers that added a 6-nt barcode for bin and replicate identification (BRBC)^70^. For every 10 OD_600_-mL of yeast used for plasmid extraction, 3 μL of plasmid template was used in a 10 μL Q5 PCR reaction (NEB).

AncSR1- and AncSR2-specific primers were mixed proportionally to background-specific cell counts (estimated from flow cytometry) to minimize amplification bias. To introduce nucleotide diversity for improved cluster identification during Illumina sequencing, eight unique forward and reverse primer pairs were used per bin and background to encode frameshift diversity and attach read 1 primer sequences in both directions. PCR conditions included 52°C annealing for 13 cycles. Reactions were then pooled by bin/replicate and purified using the Zymo DNA Clean & Concentrator Kit. In the second round, half of the first-round product was amplified with primers to add Illumina P5 and P7 adapter sequences. PCR was performed in 50 μL Q5 reactions (NEB) per 10 μL round 1 product reaction at 68.4°C annealing for 12 cycles. The final product was size-selected on a 2% agarose gel, excised, purified using the Qiagen Gel Extraction Kit, and re-purified with the Zymo DNA Clean & Concentrator Kit.

Final sequencing library concentrations were quantified by Qubit. Libraries were pooled according to the number of cells sorted per bin/replicate, and 1.8 pM dilutions were prepared according to Illumina’s standard protocol. Replicate 1 of the binned sort libraries was sequenced on a NextSeq High Output run. The remaining replicates were sequenced on a NovaSeq S1 run at the University of Chicago Genomics Facility. We used standard read primers and 86 cycles for read 1 and 80 cycles for read 2. This enabled us to bidirectionally sequence the region containing the variable RH codons and REBC.

#### Mean fluorescence estimation, data cleaning and validation

Sequencing reads were processed using a custom pipeline. We used *sickle* v1.33^71^ to filter reads based on their quality: we kept reads with a Phred score ≥ 30 and a minimum length of 79 nucleotides. We then used *PEAR* v0.9.6^72^ to merge the trimmed paired-end reads (minimum assembly length 100 nucleotides). Finally, we used Biopython toolkit v1.79^73^ to demultiplex the assembled reads by DBD background, REBC, and BRBC. We only considered reads that mapped exactly to the DBD background and allowed reads with at most one mismatch in the REBC and one in the BRBC.

The mean fluorescence for protein:RE complexes observed in the binned sort data was estimated as previously described^32^. We first estimated the proportion of cells of each complex *g* in each bin *b* (*c_g,b_*) from the proportion of reads in *b* that mapped to *g*. The mean fluorescence estimate *F_g_* for each complex was then estimated by taking the weighted mean fluorescence across bins (mean fluorescence of each bin was measured during sorting), with weights *cg*,*b*/∑*b cg*,*b*.

We applied several filtering and correction steps to reduce global measurement error and normalize fluorescence estimates between replicates. First, complexes with fewer than 27 reads per replicate were removed to ensure >95% had a standard error (SE) of ≤ 0.1 (5% of the assay range; Extended Data Fig. 2a). Second, complexes observed in only one replicate were excluded. Third, batch effects were corrected by fitting I-splines to normalize fluorescence between replicates (Extended Data Fig. 2b). Finally, SE was recalculated and complexes with SE > 0.1 were removed (Extended Data Fig. 2c). The final dataset had a mean pairwise Pearson’s *r*² = 0.55 across replicates. The poor correlation arises primarily because the vast majority of complexes are at the lower fluorescence bound, so *r*^2^ is dominated by measurement noise; for variants with fluorescence above the lower bound (roughly *F* ≥ –4.0), *r*² improved to 0.92. Altogether, we obtained fluorescence estimates for 628,732 AncSR1 and 658,475 AncSR2 variants, covering 24.6% and 25.7% of possible variants, respectively (excluding nonsense variants).

Many variants were observed at high read depth in the GFP– bin of the enrichment sort but not in the binned sort. We assigned these a null phenotype (lower-bound fluorescence) using a statistical procedure based on read depth (see Supplementary Methods), resulting in 859,171 AncSR1 and 638,762 AncSR2 protein:RE null complexes (FDR = 0.1; Extended Data Fig. 2d). This increased the total phenotyped variants to 1,487,903 in AncSR1 and 1,297,237 in AncSR2, covering 58% and 51% of all possible variants, respectively.

To evaluate the accuracy of the sort-seq fluorescence values, we measured the fluorescence of 5 isogenic variants by flow cytometry, which were also spiked into the DMS libraries prior to the binned sort. We found a high correlation between the fluorescence estimates from flow cytometry and sorting (Pearson’s *r*^2^ = 0.87, Extended Data Fig. 2e). We additionally compared the fluorescence estimates of the same variants that were contained in the DMS libraries and again observed a strong correlation with flow cytometry measurements (Pearson’s *r*^2^ = 0.97, Extended Data Fig. 2e).

To evaluate whether the REBC mutations affected fluorescence, we constructed AncSR1 and AncSR2 “mini-libraries” consisting of each of the 16 REBCs engineered into the respective wild-type protein variant. These were transformed via electroporation into the ERE or SRE reporter strain, respectively, at 1:16 the scale of the full libraries, and spiked into the full-scale libraries before sorting. The fluorescence of the mini-library variants did not differ significantly by REBC (*p* = 0.98 AncSR1, *p* = 0.99 AncSR2, one-way ANOVA), indicating that fluorescence estimates are directly comparable between libraries with different REBCs.

#### Fluorescence inference for missing complexes

To predict the fluorescence of the remaining complexes for which we did not obtain experimental estimates, we fit a generalized linear model based on reference-free analysis (RFA)^36,37^ to the experimental data. The model estimates a sigmoid function to capture the measurement bounds of the assay, plus additive and interaction effects (specific epistasis) for all amino acid states at the four variable sites in the DBD and all nucleotide states at the two variable sites in the RE. All possible intramolecular interactions up to third order amino acid interactions in the DBD and second order nucleotide interactions in the RE, and intermolecular interactions up to third order amino acid-by-second order nucleotide interactions were included. L2 regularization with 10-fold cross validation was used to reduce overfitting (Extended Data Fig. 3a; Supplementary Methods). We fit separate RFA models for each ancestral background using the *glmnet* v4.1-6 R package^74^. Model fits to the observed data were *R*^2^ = 0.96 for AncSR1 active complexes (0.31 all complexes) and *R*^2^ = 0.99 for AncSR2 active complexes (0.88 all complexes) (Extended Data Fig. 3b). These models were used to predict fluorescence values for unobserved protein-RE complexes. We also used the fitted models to correct the predictions for complexes in one of the modified strains that had systematically lower fluorescence (Supplementary Methods; Extended Data Fig. 3c, d).

#### Classification of functional complexes

We classified complexes as functional if their fluorescence was not significantly lower than the wild type complex, *i.e.* EGKA:ERE in the AncSR1 background and GSKV:SRE in the AncSR2 background. Complexes inferred as null from the enrichment sort were classified as nonfunctional. For complexes observed in the binned sort, we used a *t*-test to account for measurement error. For complexes with predicted fluorescence from the RFA models, we performed a nonparametric bootstrap test using the distribution of model residuals concatenated over the ten cross-validation fits to account for model prediction error (Supplementary Methods; Extended Data Fig. 3e). For both tests, we used a Benjamini-Hochberg FDR threshold of 0.25 to classify variants as nonfunctional if they were significantly less fluorescent than the wild type complex (Extended Data Fig. 3f). The low stringency of the FDR threshold was chosen to reduce the false positive rate for calling variants functional. The majority of complexes classified as functional in both backgrounds had fluorescence estimates obtained from the binned sort experiment (59.3% AncSR1, 75.4% AncSR2; Extended Data Fig. 3g).

#### Protein genotype networks

Following Maynard Smith’s sequence space formalism^42^, we built genotype networks consisting of all functional RH variants in each DBD background. RH genotypes are connected by an edge if they differ by a single amino acid mutation that can be produced via a single nucleotide mutation given the standard genetic code. Genotype networks for joint protein-DNA models follow a similar logic (Supplementary Methods). We used the R package *igraph* v1.5.1^75^ to build and analyze the genotype networks, and the software *gephi* v0.10.1^76^ for network visualization. To identify clusters of densely connected genotypes within the networks, we used the *cluster_edge_betweenness* function from the R *igraph* package.

#### Model of evolution on GP maps

We modeled evolution on the genotype networks as an origin-fixation process under a strong selection-weak mutation regime^77,78^ To isolate the effect of the GP map’s structure on evolution, we considered a scenario in which all functional genotypes have equal fitness, so the fixation probability is affected only by drift, and nonfunctional variants are removed by purifying selection. The relative probability *P*(*i,j*) of substitution from protein genotype *i* to genotype *j* is therefore equal to the amino acid mutation rate *µ_i,j_*, normalized over all single-step neighbors of *i* in the network. We assumed that there are no biases in the nucleotide mutation process (*e.g.* transition vs. transversion rate), so *µ_i,j_* is affected only by unequal mutational access between amino acids imposed by the genetic code. To incorporate this effect, we scaled *µ_i,j_* by the number of possible nucleotide mutations that can change any nucleotide sequence that encodes *i* to any nucleotide sequence that encodes *j:*

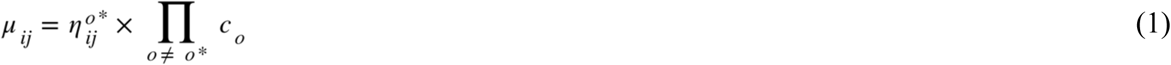

where *o* indexes the amino acid position, *o\** is the position at which the amino acid change occurs, 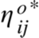 is the number of possible single nucleotide changes that can produce the state in *j* from the state in *i* at site *o*,* and *c_o_* is the number of possible codons for the invariant amino acid state at site *o*.

We used these transition probabilities to specify a discrete time Markov model for each ancestral genotype network, where each step is a single amino acid substitution. Genotypes that are more than one nucleotide change apart cannot access each other in a single time step, and the probability of staying in the same genotype across a single step in the Markov chain is also zero. We only considered functional genotypes within the main component of each network (the largest connected component). With this model, we computed the probability distribution of evolving all possible genotypes after *k* substitution steps given any specified set of starting genotypes:

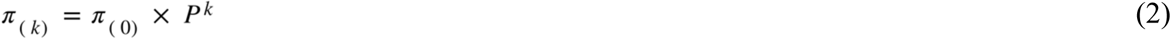

where *P* is the transition matrix with entries *P*(*i, j*), *k* > 0, and is the vector of the probability distribution of genotypes at time step *k* = 0. Setting a single element *i* of to 1 and all others to zero corresponds to evolution from a single starting genotype; setting all elements of to 1/*n*, where *n* is the number of functional genotypes in the network, averages over all possible starting genotypes. We calculated the relative probability of evolving a given specificity phenotype at time step *k* by summing over all elements of that encode that specificity and normalizing by the total probability across all specific protein genotypes.

#### Effects of background substitutions

To estimate the effect of the background substitutions between AncSR1 and AncSR2 on binding affinity, we first considered a model where the background substitutions have a universal nonspecific effect on affinity across all RH and RE genotypes. We assumed that fluorescence is proportional to the fraction of protein bound to DNA. If a complex *g* has dissociation constant *K_d_*(*g*) in the AncSR1 background, then its AncSR1 fluorescence (normalized to scale between 0 and 1) is:

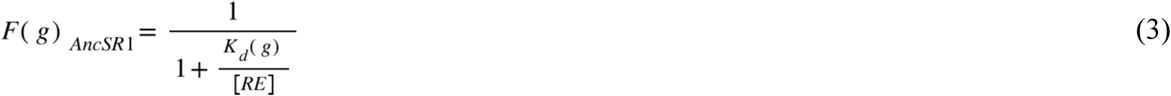

where [*RE*] is the concentration of RE. If the background substitutions scale *K_d_*(*g*) by a factor *α*, then fluorescence in the AncSR2 background is

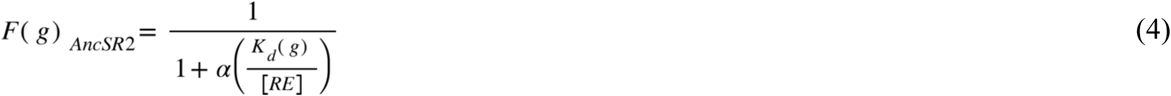

Rearranging these equations gives an expression for fluorescence in the AncSR2 background as a function of fluorescence in the AncSR1 background and *α*:

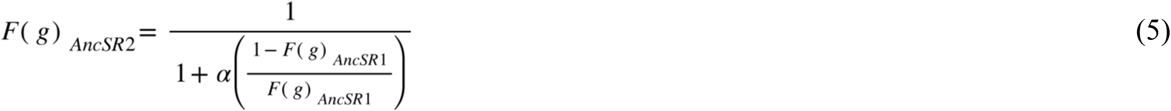

We fit this model to the AncSR1 and AncSR2 fluorescence data using orthogonal regression, which accounts for measurement error in both backgrounds. We used only complexes that had fluorescence measurements from the binned sort in both backgrounds, and whose fluorescence was significantly greater than that of nonsense variants in either background (*n* = 2,627). Fluorescence was normalized in each background to scale between the upper and lower bounds inferred from the RFA models. Confidence intervals (CI) were constructed by bootstrapping the data and refitting the model. The effect of the background substitutions was estimated to be *α* = 0.014 (95% CI: 0.010–0.014), corresponding to a 70-fold increase in affinity (95% CI: 70–99).

We next considered a model where the background substitutions have a different effect on ERE affinity than they do on other REs. We modified the model such that *α*_1_ represents the ERE-specific effect of the background substitutions and the *α*_2_ effect on the other 15 REs. We fit this model as before and obtained parameter estimates of *α*_2_ = 0.43 (95% CI: 0.19–0.76) and *α*_2_ = 0.010 (95% CI: 0.0028–0.010), corresponding to fold-increases in affinity of 2.3 (95% CI: 1.3–5.2) on ERE and 99 (95% CI: 99–361) on other REs.

## Data availability

The datasets generated and analyzed during this study are available at https://doi.org/10.5061/dryad.18931zd7m

## Code availability

Detailed scripts including how to use and analyze the data are available at www.github.com/JoeThorntonLab/RH-RE_scanning.

## Acknowledgements

We thank Yeonwoo Park and Brian Metzger for advice throughout the project and all Thornton Lab members for comments on the manuscript. This work was supported by the University of Chicago’s Research Computing Center, Cytometry Core, and Genomics Core. Funding was provided by the National Institutes of Health grants R01GM131128 (J.W.T.), R01GM121931 (J.W.T.), R35GM145336-01 (J.W.T.), T32GM007197 (J.E.J.P.), an NSF Graduate Research Fellowship (J.E.J.P.), a Rosemary Grant Award from the Society for the Study of Evolution (S.H.A.), and a Student Research Award from the American Society of Naturalists (S.H.A).

## Author contributions

All authors conceived the project; S.H.A. and J.E.J.P. performed experiments and analyzed data; all authors interpreted results and wrote the manuscript.

## Competing interests

The authors declare that they have no competing interests.

## Additional information

Supplementary information is available for this paper. Correspondence and material requests should be addressed to J.W.T.

---

**Extended Data Figure 1.**
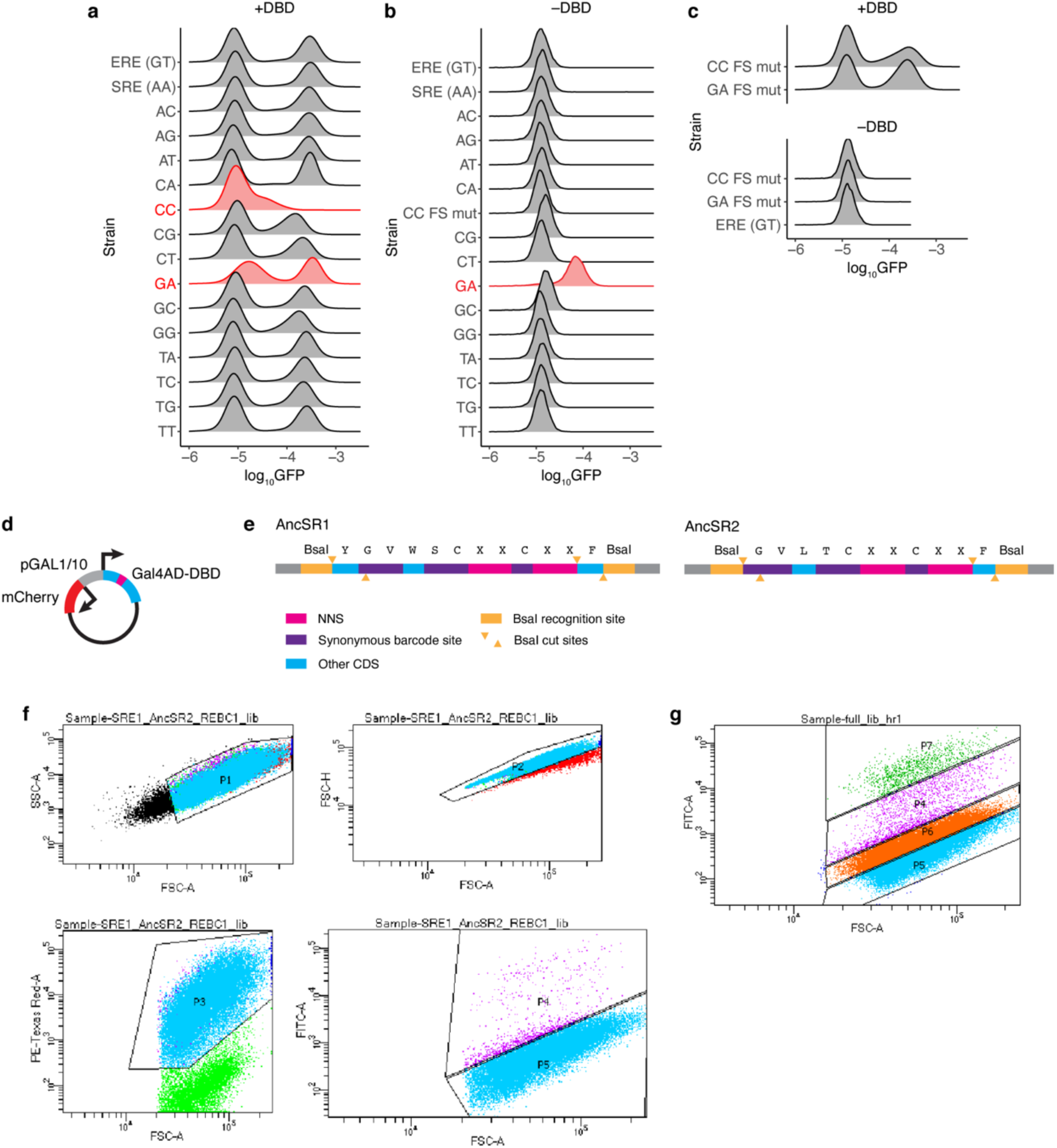
DBD library construction and sorting. **a-c**, Validation of the RE reporter strains. GFP fluorescence was measured by flow cytometry in each strain in the presence (+DBD) or absence (–DBD) of a universally high-affinity DBD variant (AncSR1+GGKA+11P, ^28^). In each row, the left peak corresponds to autofluorescence from cells that do not express GFP, either due to lack of DBD binding or loss of the DBD expression plasmid; the right peak corresponds to cells that are expressing GFP in response to DBD-RE binding. “FS mut” denotes strains with mutations in the flank/spacer regions of the RE that correct anomalous expression patterns shown in red (see Supplementary Methods). Red strains were not used in the final DMS experiment. Experiments were conducted on the same day within each panel. Fluorescence is shown in the presence of high-affinity DBD (panel a), in the absence of DBD expression plasmid (panel b), and for the CC and GA FS mut strains, with the ERE strain included as a negative control (panel c). **d-e**, Design of DBD library oligos. NNS codons (pink) were used to generate all possible combinations of amino acid mutations at the four RH scanning sites (marked as X in the amino acid sequence). For each background (AncSR1, left; AncSR2, right), we synthesized 16 libraries, each with a unique set of synonymous barcode mutations at five codons (purple, Supplementary Table 1), which allows each to be associated with one RE strain. BsaI sites (orange) were used for Golden Gate assembly into the pDBD2.1 backbone. **f-g**, Sorting gates used for DMS. **f**, Enrichment sort gates. Homogeneous single cells were first selected by gating on FSC-A vs. SSC-A and FSC-A vs. FSC-H (top). Plasmid retention was then selected for by gating on mCherry expression (PE-Texas Red-A, bottom left). Finally, cells were sorted into GFP+ (P4) and GFP– (P5) populations (bottom right). The boundary between the GFP+ and GFP– gates was drawn to have a slope of 1.5 on a log-FSC-A vs. log-GFP (FITC) scale so that populations were sorted by GFP expression relative to cell volume. **g**, Binned sort gates. Gates P1–P3 were drawn as in **C**. Cells were then sorted into four GFP bins, which were drawn to have roughly equal heights (P5–P7). The boundaries between GFP gates were again drawn to have a log-log slope of 1.5.

**Extended Data Figure 2.**
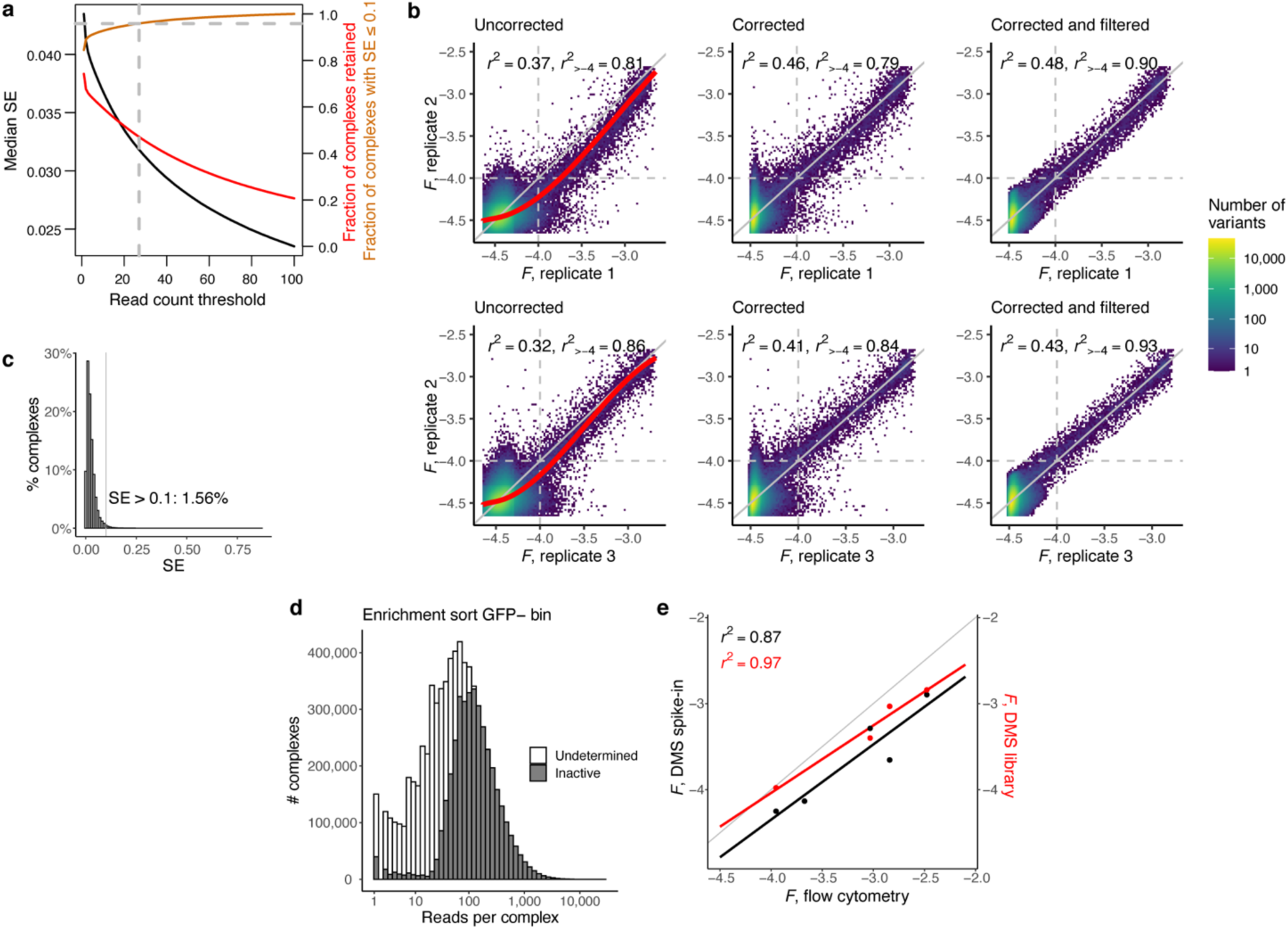
DMS data cleaning. **a**, Curves show characteristics of the binned sort dataset as a function of the read count threshold used to retain protein-RE complexes for further analysis (*x*-axis). Black, standard error of *F* (SE, left axis); red, complexes retained, expressed as a fraction of the number of complexes in the binned sort (right axis); gold, fraction of complexes retained that have SE ≤ 0.1 (right axis). We used a read count threshold of 27 (vertical dashed line), at which ≥95% of complexes have SE ≤ 0.1 (horizontal dashed line). **b**, Correcting and filtering estimates of *F* from the binned sort. Left, correlation in *F* between replicates before correction. Pearson’s *r*^2^ is shown for all complexes, and for the subset of complexes with *F* > –4 in both replicates, which roughly corresponds to the boundary between active and inactive complexes (gray dotted lines). Red curves, I-splines fit using complexes with SE of *F* < 0.1. Center, correlation in *F* between replicates after correcting using the I-spline transforms. Right, correlation in *F* between replicates after filtering corrected variants for SE ≤ 0.1. **c**, Distribution of SE across all complexes in the binned sort after the I-spline correction. Complexes with SE > 0.1 were discarded. **d**, Read count distribution for complexes sequenced in the enrichment sort GFP– bin. Complexes were inferred to be inactive (gray) if they were not observed in the binned sort, but had high enough inferred cell count in the enrichment sort to have been detectable in the binned sort had they been at least minimally fluorescent (see Supplementary Methods). **e**, Correlations between estimates of *F* from flow cytometry (*x*-axis) and DMS (*y*-axes). Left *y*-axis (black points) shows estimates from isogenic strains that were spiked into the DMS libraries prior to the binned sort. Right *y*-axis (red points) shows estimates from complexes that were encoded in the DMS libraries.

**Extended Data Figure 3.**
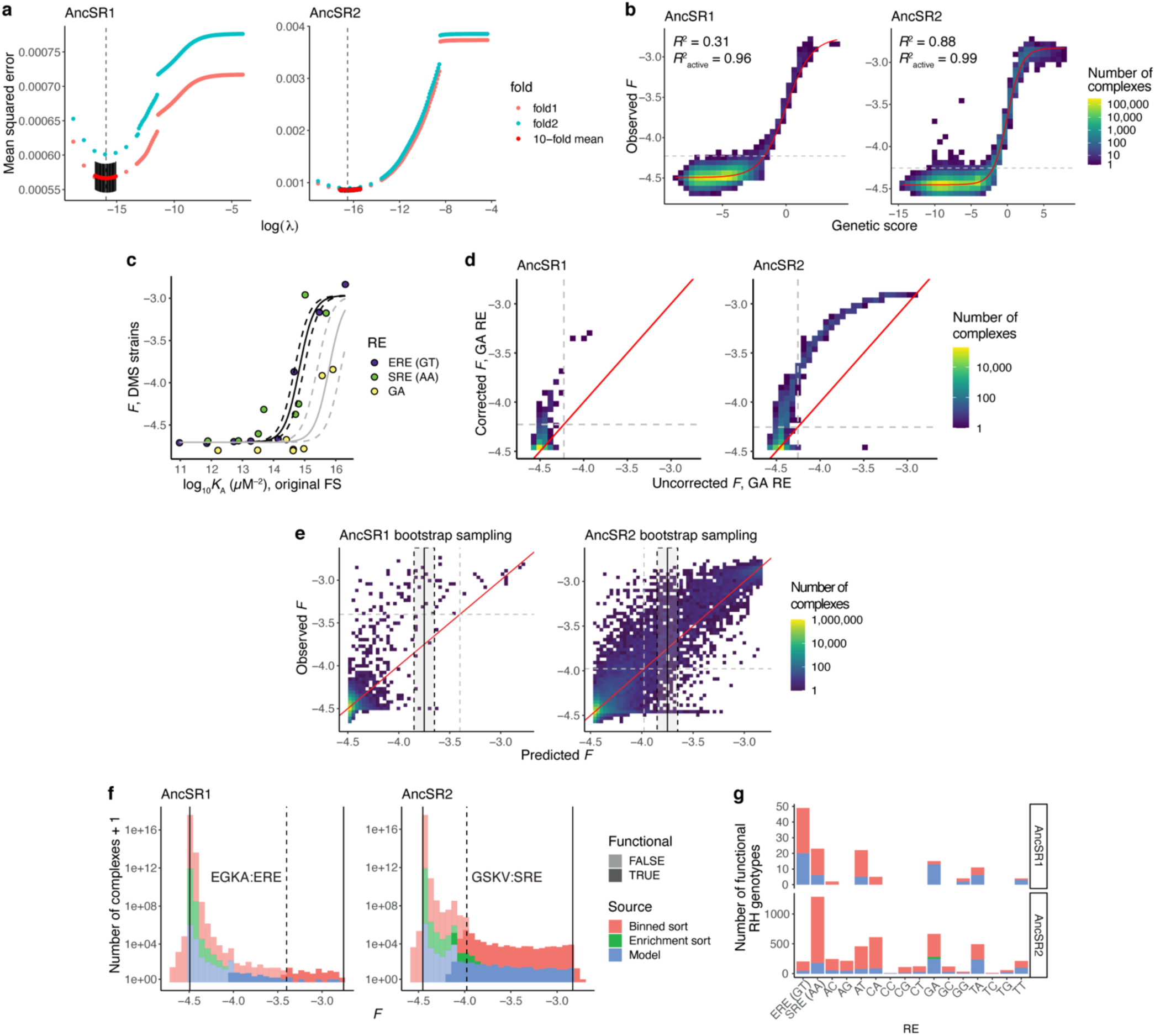
Fluorescence imputation, GA fluorescence correction, and functional genotype classification. **a**, **b**, A generalized linear model that predicts the fluorescence of each protein-RE complex from its sequence was fit to the data for each background, using L2 regularization to address overfitting. **a**, Ten-fold cross-validation (CV) was used to identify the optimal L2 penalty parameter (λ). Red and black, mean and SE of the out-of-sample mean squared error (MSE) across the 10 folds. Initial range finding was performed using two folds (pink and cyan). Vertical line, λ that minimizes mean MSE. **b**, Genetic score versus observed *F* for the regularized RFA models. Red line, best-fit nonspecific epistasis function. For display, the distribution was discretized; colors show the number of variants in the interval defined by each square. Coefficient of determination (*R*^2^) is reported for all complexes and for the subset of active complexes (above the gray line). **c**, **d**, Fluorescence correction for the GA strain. **c**, Affinity (*K*_A_) versus *F* for a panel of DBD variants measured on ERE, SRE, and GA. Affinities, measured by fluorescence anisotropy on the three REs, all with the original flank/spacer sequence, were previously reported^28,33^. *F* was measured by flow cytometry in the RE strains that were used for DMS, of which the ERE and SRE strains had the original flank/spacer sequence, and the GA strain had a mutated flank/spacer sequence (see Supplementary Methods, Extended Data Fig. 1c–e). Curves, best-fit sigmoidal function. The same midpoint parameter was used for ERE and SRE (black); that for GA was independently estimated (gray). Dashed lines, sigmoidal functions using 95% confidence intervals on the midpoints. **d**, GA fluorescence correction based on the affinity effect estimated in **c**. Plots show *F* before and after the correction. Dashed gray lines, mean boundary between active and null variants. Red line, *y* = *x*. **e**, Bootstrap sampling strategy for classifying functional complexes with model-inferred fluorescence. Plots show concatenated out-of-sample predictions versus observed *F* across all 10 CV models. Bootstrap-sampled residuals from the interval within ±0.1 units of a complex’s predicted *F* were used to test whether a variant with model-inferred *F* was not significantly worse than the wild-type complex (dashed gray lines). An example for a complex with inferred *F* = –3.75 (solid black line) is shown, with the bootstrap interval shown as a shaded rectangle. Solid red line, *y* = *x*. **f**, Distribution of *F* across all 2,560,000 complexes in each DBD background. Solid vertical lines, upper and lower bounds of fluorescence inferred from the RFA models; dashed vertical lines, fluorescence of wild type complex (EGKA: ERE for AncSR1 and GSKV:SRE for AncSR2). Colors indicate the source from which *F* was estimated. Darker colors show functional variants, lighter colors nonfunctional. All “enrichment sort” complexes were assigned to the lower bound of fluorescence, except for GA RE variants whose fluorescence was corrected upward (**d**). Some model-predicted variants in the AncSR1 background have predicted *F* below the reference but are classified as functional, because the bootstrap test accounts for the AncSR1 RFA model’s tendency to under-predict fluorescence (**e**, left). **g**, Bars show the number of functional RH variants per RE per DBD background, colored by source of *F* estimate as in **f**.

**Extended Data Figure 4.**
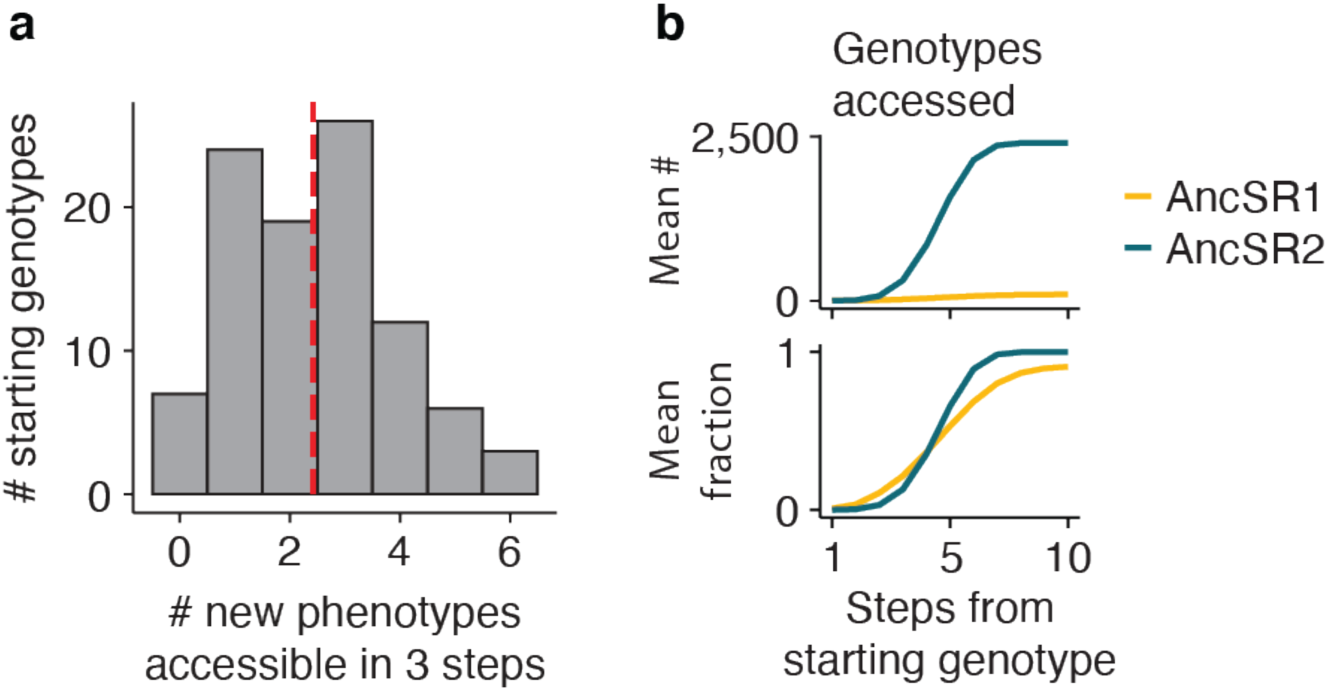
Additional network analyses. **a**, Accessible new phenotypes after 3 substitution steps in the AncSR1 network. Bars show the distribution over every starting genotype in the AncSR1 main component. Dashed line, mean. **b**, Number (top) and fraction (bottom) of genotypes in each network accessible as a function of the length of evolutionary trajectories. Lines show the mean across all starting genotypes in each network. Gold, AncSR1 network; teal, AncSR2 network.

**Extended Data Figure 5.**
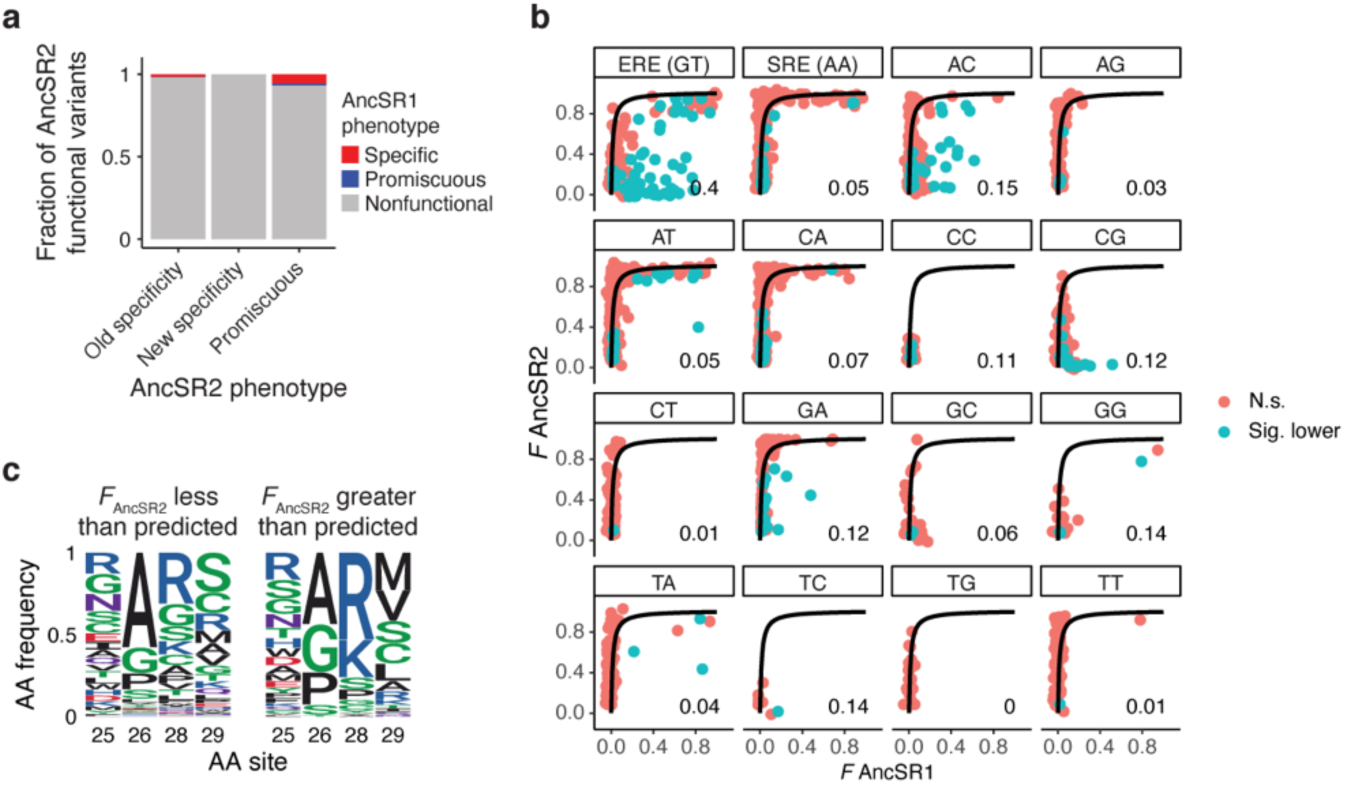
Additional analyses for effects of background substitutions on DBD-RE affinity. **a**, Changes in phenotype across the AncSR1-to-AncSR2 transition. Bars represent the set of protein variants in AncSR2 that have different classes of phenotypes: specificity phenotypes that were encoded in the AncSR1 map (old specificity), specificity phenotypes not encoded in the AncSR1 map (new specificity), or promiscuous in AncSR2. Colored sections show the fraction of variants in each class whose functional category in the AncSR1 background was specific, promiscuous, or nonfunctional. **b**, Plots are the same as in Fig. 6a, but split into panels by RE. Blue points, protein-DNA complexes with significantly lower fluorescence in the AncSR2 background than predicted by the model; red, all other variants. Numbers at the bottom-right of each panel show the fraction of plotted variants with significantly lower than expected AncSR2 fluorescence. **c**, Amino acid frequencies at the RH variable sites among all complexes that are significantly more (left) or less (right) fluorescent in the AncSR2 background than predicted by the ERE-specific model in Fig. 6e. To test for significance in **b** and **c**, we tested whether their Bonferroni-corrected 95% CI of fluorescence was outside of the 95% CI of the model in both the AncSR1 and AncSR2 backgrounds.

**Extended Data Figure 6.**
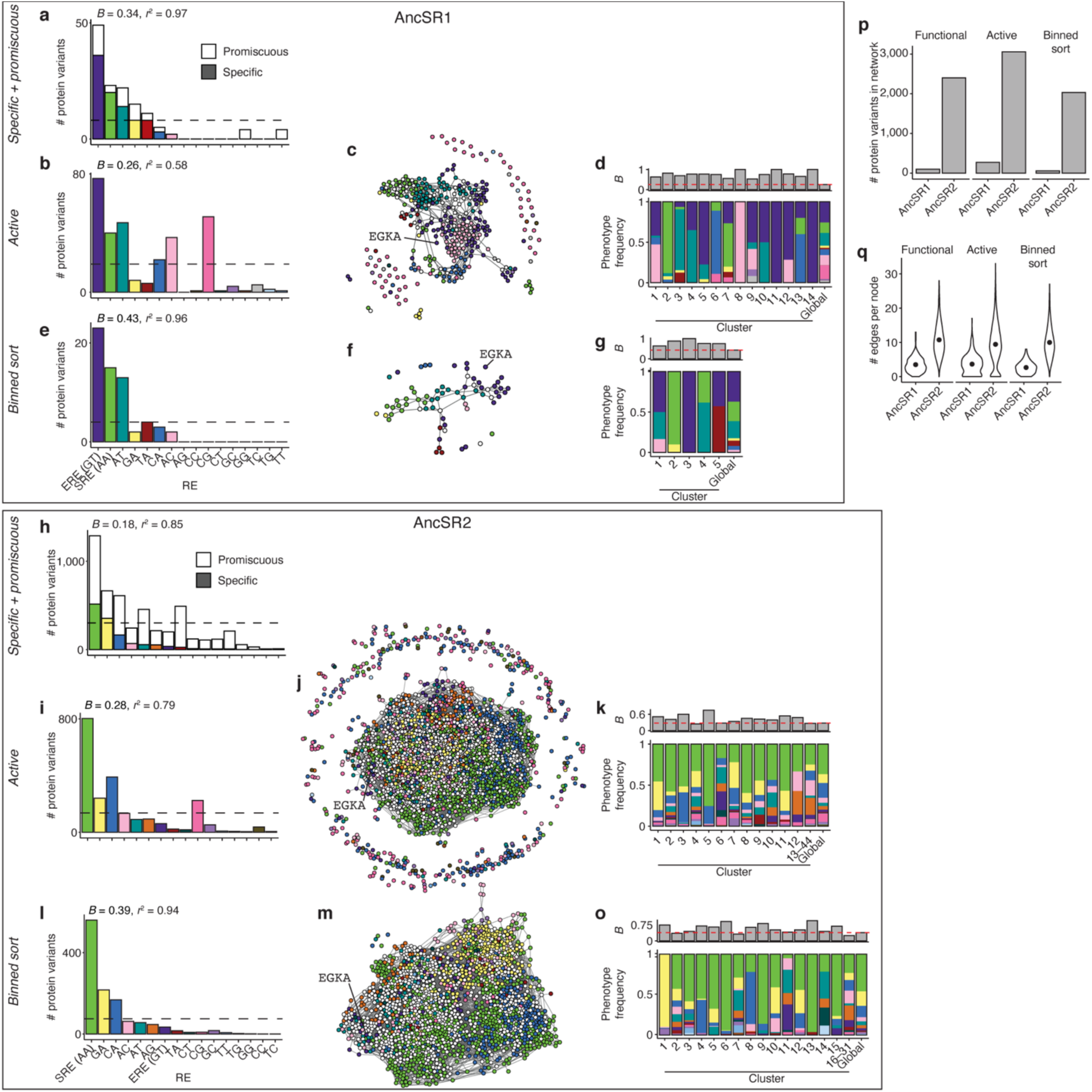
Robustness to alternative phenotype assignment methods. **a**, Global production distribution in the AncSR1 background, counting variants that bind specifically (colored bars) and promiscuously (white bars) to each RE. Dashed line shows the expected frequencies if the production distribution were isotropic. Nonuniformity, *B*, of the distribution and *r*^2^ to the production distribution for specific variants (Fig. 2a) are reported. **b**, Same as in **a**, with phenotypes calculated using data from variants with fluorescence significantly higher than that of nonsense variants (active variants). **c**, Sequence space network for AncSR1 active variants. **d**, Bottom: Frequencies of specificity phenotypes within each genotype cluster in the AncSR1 active variant networks; the global production distribution is shown for comparison. Top: *B* in each cluster. Red line, *B* of global production distribution. **e–g**, Same as in **b–d**, but with phenotypes calculated using only data from the binned sort experiment; protein-DNA complexes without experimental fluorescence measurements were assumed to have null fluorescence. **h–o**, Same as in **a–g**, but for the AncSR2 background. Note that the active variant datasets are likely to be enriched for false positives due to misclassification of variants whose fluorescence is by chance slightly higher than the nonsense variant distribution. This may explain the high frequency of variants that do not share any mutational connections to other active variants. It may also explain the high frequency of CG-specific variants compared to the original classification scheme, since the CG yeast strain has a slightly higher null fluorescence level than most other strains (Extended Data Fig. 1c, d) and most CG-specific variants are unconnected in the active variant genotype networks. **p**, Number of protein variants in each network under different methods of phenotype assignment. “Functional” indicates the original method used in the main text; note that this yields the same number of protein variants as the “specific + promiscuous” method. **q**, Number of edges per node in each network, with the original phenotype classification method (functional) shown for comparison.

**Extended Data Figure 7.**
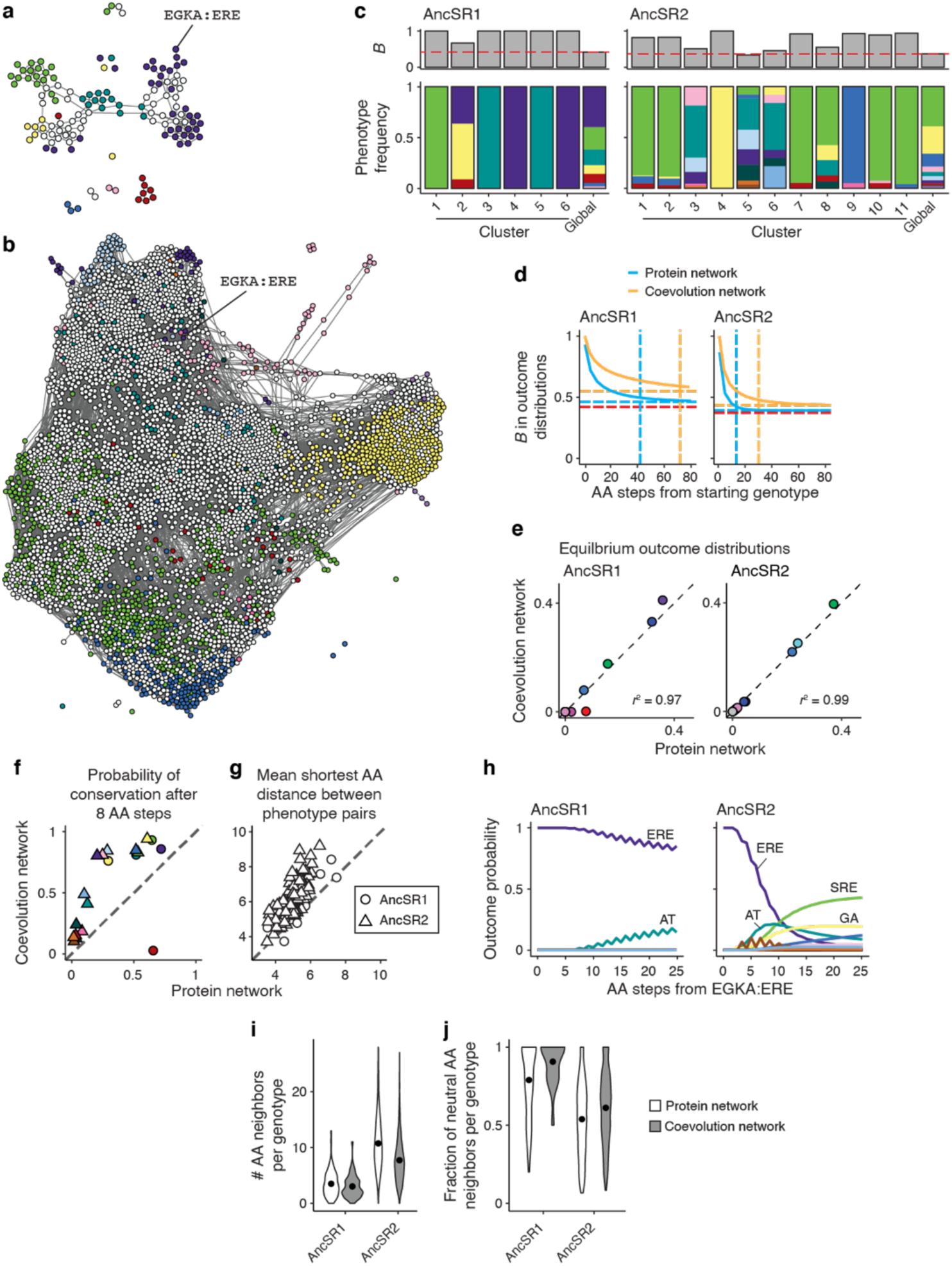
Robustness to model of evolution using joint protein-DNA networks. **a**, AncSR1 protein-DNA coevolution network. Nodes represent functional protein-RE complexes, colored by the RE specificity of the protein genotype; colors are as in Fig. 2b and 4b. Promiscuous protein genotypes are represented by multiple nodes, one for each RE it binds. Edges connect complexes that can be interconverted by a single nucleotide change in the RE or the coding sequence of the protein. **b**, AncSR2 protein-DNA coevolution network. **c**, Bottom: Frequencies of specificity phenotypes within each genotype cluster in the AncSR1 (left) and AncSR2 (right) coevolution networks; the global production distribution (right-most column) is shown for comparison. Top: Nonuniformity of phenotype distribution (*B*) within each cluster. Red line, *B* of global production distribution. **d**, *B* in evolutionary outcomes as a function of the length of evolutionary trajectories. Solid curves, mean *B* across starting genotypes in the protein (cyan) or coevolution (orange) networks. Dashed horizontal lines, *B* of the equilibrium outcome distribution in each network; dashed horizontal red line, B of the frequency distribution of phenotypes in the network. Vertical dashed lines show the number of substitutions required for mean *B* to reach within 0.05 units of the equilibrium value within each type of network. The equilibrium distributions are more anisotropic in the coevolution networks, and require more amino acid substitutions to be reached, because changes in protein genotype must occur between variants that can bind to the same RE sequence. **e**, Comparison between equilibrium outcome distributions of the protein-only evolution and protein-DNA coevolution networks in each AncSR1 (left) and AncSR2 (right) backgrounds. Pearson’s *r*^2^ between the two distributions are shown. Dashed line, *y* = *x*. **f**, Probability of conservation of each phenotype after 8 amino acid substitution steps in the protein vs. coevolution networks. **g**, Mean shortest amino acid distance between all possible pairs of phenotypes in the coevolution vs. protein networks, calculated as in Fig. 2g. Circles, AncSR1 networks, triangles, AncSR2 networks. Dashed line, y = x. **h**, Probability of evolving each specificity phenotype as a function of the number of amino acid substitutions away from EGKA:ERE in the AncSR1 (left) and AncSR2 (right) coevolution networks. In both backgrounds, conservation is more likely at short trajectory lengths than in the corresponding protein networks (Fig. 3g, 5f), but the relative likelihood of achieving each phenotypic outcome is similar. **i**, Distribution of the number of neighbors per genotype with distinct RH sequences in each type of network. Dots, means of distributions. **j**, Distribution of the fraction of neutral neighbors per node with distinct RH genotypes in each network. Dots, means of distributions.

**Extended Data Figure 8.**
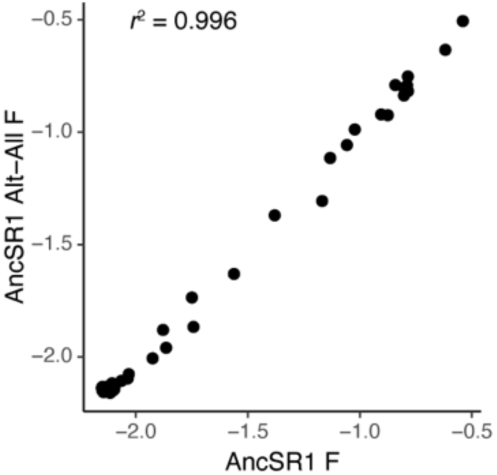
Robustness of RH mutation effects to uncertainty in ancestral reconstruction. Effects on ERE binding of all possible single amino acid mutations at the four variable RH sites in the background of the maximum *a posteriori* (MAP) wild type AncSR1 protein (*x*-axis), and in the background of the AltAll wild type AncSR1 protein, which has the second-most likely amino acid state at all sites at which the posterior probability of the MAP state is less than 0.8 (*y*-axis)^32^. Pearson’s *r*^2^ is shown.

**Extended Data Figure 9.**
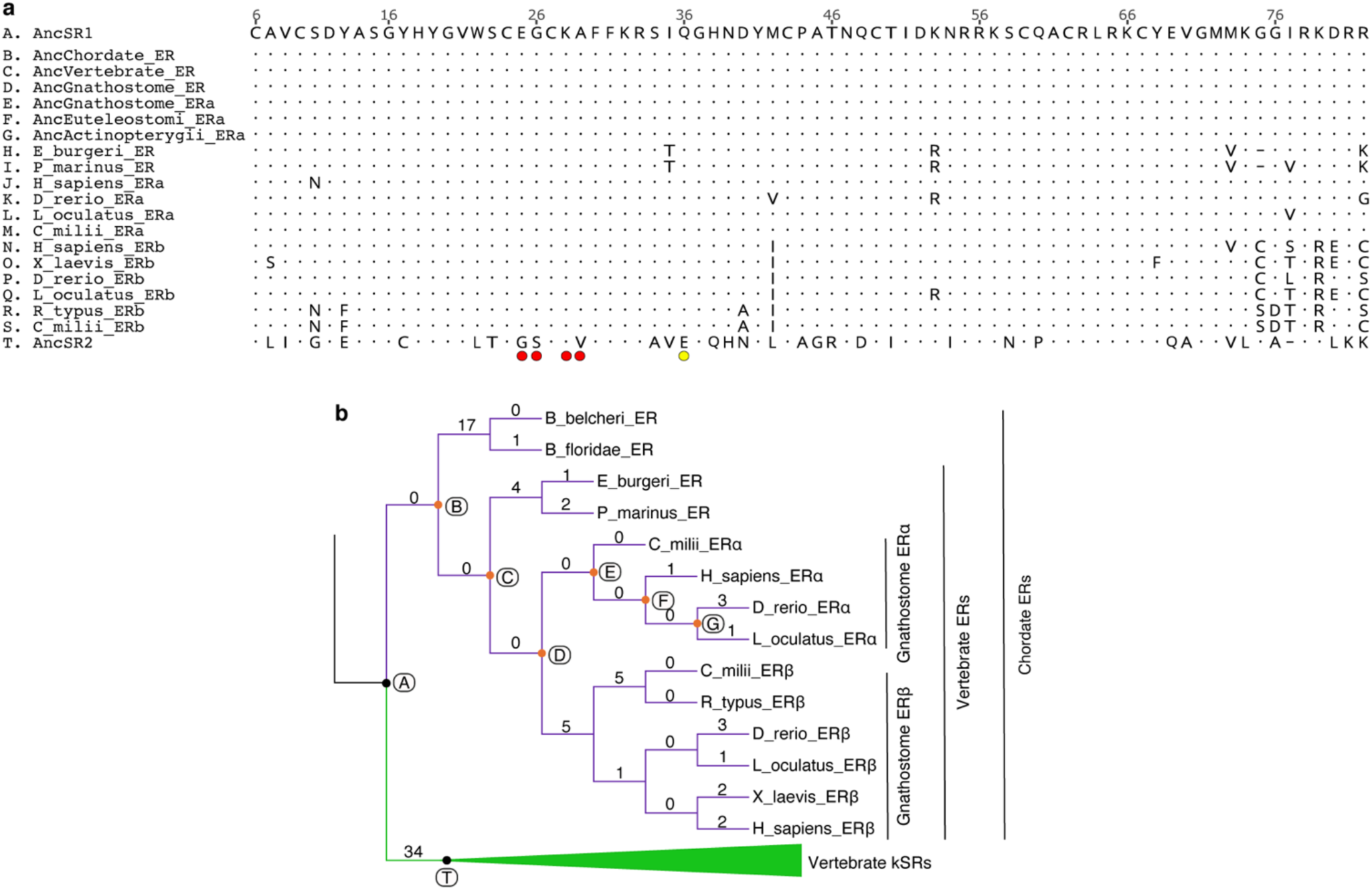
Amino acid changes along the SR phylogeny. **a**, Amino acid alignment of extant vertebrate ERs and the MAP protein sequences for key ancestral nodes in the SR phylogeny^32^. The AncSR1 sequence is used as the reference to indicate amino acid changes; dots, same amino acid state as that in AncSR1; dashes, gaps; red circles, variable sites in DMS experiment; yellow circle, historical substitution (q36E) that likely contributed to the shift in the direction of anisotropy away from ERE. **b**, Cladogram of SRs showing the number of substitutions that occurred along each branch. Letters, nodes shown in alignment in **a**; black nodes, AncSR1 and AncSR2; orange nodes, ancestral ER sequences identical to AncSR1. Branches and clades are colored according to their DNA specificity phenotype: purple, ERE-specificity; green, SRE-specificity.

## Notes

### Competing Interest Statement

The authors have declared no competing interest.

### Summary of Updates

Modified the text and title to more precisely describe the experiments and the rationale for them.

