## Supplementary Information for "The structure of an ancient genotype-phenotype map shaped the functional evolution of a protein family"

**Supplementary Information for "Biases in an ancient genotype-phenotype map drove functional evolution in the steroid hormone receptor family"**

Santiago Herrera-Álvarez†, Jaeda E. J. Patton†, Joseph W. Thornton*

†Co-first authors

### CONTENTS

- ***Supplementary Methods.*** Methodological details not described in the main Methods, including protocols for library assembly and high-throughput yeast transformation.

- ***Supplementary Table 1: Synonymous RE barcodes (REBCs).*** Sequences of synonymous mutations used to barcode the DBD libraries to associate them with RE strains. NNS, variable sites in the DMS experiment.

- ***Supplementary Table 2: Library transformation and enrichment sort statistics.*** Library transformation yields, glycerol stock recovery rates, and number of cells sorted for enrichment sorts.

- ***Supplementary Table 3: Binned sort statistics.*** Glycerol stock recovery rates and number of cells sorted for binned sorts.

- ***Supplementary Table 4: Binned sort sequencing statistics.*** Estimated number of reads per cell across libraries, bins, and replicates. Libraries for AncSR1 REBC 9–14 and REBC 15–16 had very low and high sequencing depths, respectively, in replicate 1, so we repeated the enrichment sort for these libraries and used them for bins sort replicates 2–4.

- ***References for Supplementary Materials.***

### SUPPELEMENTARY METHODS

*Dynamic range correction for RE reporter strains*

14 of 16 strains showed DBD-dependent fluorescence across a similar dynamic range, with two fluorescence peaks in the presence of DBD—a GFP-negative peak (GFP–) corresponding to autofluorescence from cells that had spontaneously lost the DBD plasmid, and a GFP-positive peak (GFP+) from cells that retained the plasmid and expressed GFP—and a single GFP– peak in the absence of DBD (Extended Data Fig. 1a, b). With the CC RE strain, however, the GFP+ peak was absent (Extended Data Fig. 1a); with the GA RE, the GFP– peak was right-shifted, indicating high basal fluorescence even in the absence of DBD (Extended Data Fig. 1a, b). We repeated the transformation and obtained the same results. We hypothesized that inserting the CC and GA RE sites may have introduced cryptic yeast TF binding sites or caused chromatin remodeling that resulted in constitutive GFP repression and activation, respectively. The flank and spacer (FS) sequences surrounding the RE half sites do not affect SR binding specificity [1–3], so we reasoned that introducing mutations into these regions might preserve the ability of the strains to act as reporters for RE binding while disrupting any recognition sites for endogenous yeast proteins. We experimentally identified a set of FS mutations for the CC strain (aacAGCCCAaaaTGGGCTgtt) and another for the GA strain (ccaAGGACAatcTGTCCTtgg) that result in DBD-dependent GFP expression similar to the other RE strains (Extended Data Fig. 1c). We therefore used these two FS-modified strains along with the original strains for the remaining 14 RE variants for DMS experiments.

*Details on replicate sorting, sequencing and processing*

The binned sort was performed to yield 3 replicates per library. We sequenced and processed replicate 1 of the binned sort libraries before the other two replicates to assess data quality (see below). Analysis of this replicate revealed that the batch containing libraries with REBC 9-16 in the AncSR1 background contained inconsistent read counts—we expected about 1 read/cell, but these libraries contained very low or very high estimates of reads per cell (Supplementary Table 4). To fix this, we repeated the sorting procedure (enrichment and binned sorting) and used these data (replicate 4) to replace replicate 1 for these eight libraries (Supplementary Table 4).

Replicate 1 of the binned sort libraries was sequenced on a NextSeq High Output run. The remaining replicates (2, 3 and 4) were sequenced on a NovaSeq S1 run at the University of Chicago Genomics Facility. We obtained $1.27 \times 10^8$ and $2.1 \times 10^9$ read pairs for the NextSeq and NovaSeq runs, respectively. These encompassed the 32 experimental libraries, the isogenic controls, and the mini-libraries. We used *sickle v1.33*[4] to filter sequencing reads based on their quality: we kept reads with a Phred score ≥ 30 and a minimum length of 79 nucleotides. This procedure resulted in a cleaned dataset of $1.2 \times 10^8$ and $1.85 \times 10^9$ read pairs for the NextSeq and NovaSeq runs, respectively. We used *PEAR v0.9.6*[5] to merge the trimmed paired-end reads (minimum assembly length 100 nucleotides), resulting in $1.18 \times 10^8$ (>98%) and $1.75 \times 10^9$ (>94%) assembled reads, respectively.

We used Biopython toolkit *v1.79*[6] to demultiplex the assembled reads by DBD background, REBC, and BRBC. We only considered reads that mapped exactly to the DBD background and REBC, and allowed reads with at most one mismatch in the BRBC. We successfully mapped $1.01 \times 10^8$ (>85%) of the assembled NextSeq reads, and $1.65 \times 10^9$ (>94%) of the assembled

NovaSeq reads. For downstream analyses, we combined the mapped reads from NextSeq and NovaSeq.

We corrected the fluorescence estimates for batch effects between replicates 1-3. However, fluorescence estimates from replicate 4 were not corrected for batch effects due to the low number of active variants in the libraries.

*Statistical classification of null variants from enrichment sort GFP– bin*

In addition to quantitative fluorescence estimates from the binned sort dataset, we reasoned that we could assign variants a null phenotype (*i.e.* baseline-level fluorescence) if they were observed in the enrichment sort GFP– bin but were not observed in the binned sort data, since this implies that they did not express GFP during the enrichment sort. A complication is that variants observed at low frequency in the enrichment sort GFP– bin may simply have not been sorted to sufficient depth to be detected in the binned sort; these variants cannot be confidently classified as null. We therefore set out to test whether variants were sorted to sufficient depth in the enrichment sort to be confidently assigned a null phenotype if they did not appear in the binned sort dataset.

We reasoned that if a variant was sorted to a high enough depth in the enrichment sort to be detectable as fluorescent in the binned sort, then if it did not appear in the binned sort it should be classified as null. We therefore estimated the probability $p_m$ of failing to detect a variant $m$ as significantly fluorescent in the binned sort data; we take this to be the probability of capturing fewer than $k$ cells of variant $m$ in the enrichment sort GFP+ bin, where $k$ is the minimum number of enrichment sort GFP+ cells required to detect a fluorescent variant in the binned sort. This can be calculated as a binomial sampling probability:

$$p_m = Pr\left(c_m^{d+} < k\right) = F_{Binom}\left(k; c_m^d, f\right) \tag{3}$$

where $c_m^{d+}$ is the number of cells of variant $m$ sorted into the enrichment sort GFP+ bin, $c_m^d$ is the total number of cells of variant $m$ sorted in the enrichment sort, $f$ is the minimum fraction of cells in the enrichment sort GFP+ bin for a variant to be detected as fluorescent in the binned sort, and $F_{Binom}$ is the binomial cumulative distribution function. If $p_m$ is low then it is likely that variant $m$ was sorted to sufficiently high depth in the enrichment sort to be detected as fluorescent in the binned sort; $p_m$ can therefore be used as a $p$-value for classifying variants as null if they are not observed in the binned sort.

We first estimated $c_m^d$ for variants observed in the enrichment sort GFP– sequencing data. To do this, we estimated both the number of cells sorted into the enrichment sort GFP– bin, $c_m^{d-}$, and the number of cells sorted into the enrichment sort GFP+ bin, $c_m^{d+}$. For variant $m$ in library $l$, we took $c_m^{d-}$ to be the fraction of enrichment sort GFP– reads from library $l$ that map to variant $m$, multiplied by the total number of GFP– cells sorted for library $l$:

$$c_m^{d-} = \frac{r_m^{d-}}{r_l^{d-}} \times c_l^{d-} \tag{4}$$

where $r_m^{d-}$ is the number of enrichment sort GFP– reads for variant $m$, $r_l^{d-}$ is the total number of enrichment sort GFP– reads for library $l$, and $c_l^{d-}$ is the number of GFP– cells sorted for library $l$ (Supplementary Table 2). Since we did not sequence the enrichment sort GFP+ bin directly, to estimate $c_m^{d+}$ we assumed that the fraction of cells of variant *m* in library *l* in the binned sort data was proportional to the fraction of cells of variant *m* in library *l* in the enrichment sort GFP+ bin:

$$c_m^{d+} = \frac{c_m^b}{c_l^b} \times c_l^{d+} \tag{5}$$

where $c_m^b$ is the number of inferred binned sort cells for variant *m* (summed across all sort bins), $c_l^b$ is the total number of inferred binned sort cells for all variants in library *l* (summed across all sort bins), and $c_l^{d+}$ is the number of GFP+ cells sorted for library *l* in the enrichment sort (Supplementary Table 2). The sum of $c_m^{d-}$ and $c_m^{d+}$ is our estimate of $c_m^d$.

Next, we estimated the minimum number ($k$) and fraction ($f$) of GFP+ cells in the enrichment sort required for a variant to be detected as fluorescent in the binned sort. Variants with lower fluorescence are less likely to be detected in the binned sort since they have a lower fraction of GFP+ cells, so we set out to define a class of minimally fluorescent variants with which to estimate of $k$ and $f$. To do this, we classified variants observed in the binned sort as active (*i.e.,* significantly more fluorescent than null) or null by computing the fraction of nonsense variants of similar read depth and protein background with higher fluorescence than the test variant; this was used as a $p$-value for classifying variants as active (FDR = 0.1). Minimally fluorescent variants were defined as those with fluorescence within ±0.05 units of that of the least-fluorescent active variant. We then used the median $c_m^{d+}$ of minimally fluorescent variants, taken as a weighted average across read depth bins, to obtain an estimate of $k = 9$; the median number of cells in binned sort bins 3 and 4 (roughly equivalent to the enrichment sort GFP+ population) for minimally fluorescent variants, averaged across read depth bins, was calculated to obtain an estimate of $f = 0.23$.

Using Equation 1 from the main Methods section with our estimated parameter values, we were able to classify an additional 859,171 AncSR1 and 638,762 AncSR2 variants as null (FDR = 0.1; Extended Data Fig. 2d). This brought the total number of phenotyped variants to 1,487,903 in AncSR1 and 1,297,237 in AncSR2, corresponding to 58% and 51% of all possible variants, respectively.

*Generalized linear model to predict fluorescence of missing variants*

A remaining 42% of AncSR1 and 49% of AncSR2 variants were either unobserved or had insufficient read depth to be confidently assigned a phenotype. To predict the fluorescence of these variants, we used a type of generalized linear modeling approach called reference-free analysis (RFA)[7,8] to predict fluorescence based on the effects of sequence states inferred from empirically phenotyped variants. RFA is an unbiased method for inferring the phenotypic effects of genetic states and their interactions (*i.e.,* epistasis) in a combinatorial genotype space. Each genotype $g$ of length $n$ is represented as a vector of genetic states at each site $\left(g_1, g_2, \ldots, g_n\right)^T$. RFA relates the genetic states in $g$ to a latent phenotype $s$, also referred to as the genetic score, through a linear combination of main and epistatic effects:

$$s(g) = e_0 + \sum_i^n e_i(g_i) + \sum_{i<j}^n e_{i,j}(g_i, g_j) + \ldots + \varepsilon \tag{6}$$

Here, $e_0$ is the global mean genetic score across all variants, $e_i(g_i)$ is the first-order (average) effect of state $g_i$ at site $i$, and $e_{i,j}(g_i,g_j)$ is the pairwise epistatic effect of states $g_i$ and $g_j$ at sites $i$

and *j*; the model can be extended to include higher order epistatic interactions between sites. The genetic score is related to phenotype through a sigmoid link function:

$$F(g) = L + \frac{U - L}{1 + e^{-s(g)}} + \varepsilon \quad (7)$$

where $F(g)$ is the empirically measured phenotype of $g$, $L$ and $U$ are global parameters representing lower and upper phenotype bounds; and $\varepsilon$ is experimental noise, assumed to be normally distributed. The sigmoid link function accounts for nonspecific epistasis arising from biological and/or experimental bounds on the dynamic range of $F$.

We estimated separate RFA models for each ancestral DBD background. The model estimates the effects of all amino acid states at the 4 variable sites in the DBD and all nucleotide states at the 2 variable sites in the RE; it contains intramolecular interactions up to 3rd order amino acid interactions in the DBD and 2nd order nucleotide interactions in the RE, and intermolecular interactions up to 3rd order DBD-by-2nd order RE interactions. All variants with empirical fluorescence estimates from either the binned or enrichment sorts were used in training; for variants classified as null from the enrichment sorts, we used the mean fluorescence of nonsense variants from the same background. Models were fit in R using *glmnet* v4.1-6[9]. Because *glmnet* does not allow for estimation of parameters in the link function, we first fit an unregularized RFA model with up to 2nd order DBD-by-2nd order RE effects using nonlinear least squares regression to estimate the global $U$ and $L$ parameters. We then used these estimates to specify a link function for *glmnet*, which we used to fit the full model using L2-regularized regression. 10-fold cross validation (CV) was used to select the L2 penalty that minimized prediction error (Extended Data Fig. 3a). Our final models fit the data for active variants with $R^2 = 0.96$ (AncSR1) for and $R^2 = 0.99$ (AncSR2); for all variants, $R^2 = 0.31$ (AncSR1) and $R^2 = 0.88$ (AncSR2), because most variation in fluorescence for null variants is caused by measurement error (Extended Data Fig. 3b, c).

We used the RFA models to predict fluorescence for the missing variants in our dataset and classified these variants as null or active. The final dataset with combined empirical and predicted fluorescence estimates for AncSR1 contained 460 active protein variants, of which 114 (25%) were from model predictions; for AncSR2, there were 7,601 active variants, of which 1,838 (24%) were from model predictions.

We used the model to correct for the observation that the GA strain has systematically lower fluorescence than expected given previous measurements of affinity for this RE and a panel of protein variants (Extended Data Fig. 3c)[10], presumably because the FS mutations we introduced reduce affinity (see *Dynamic range correction for RE reporter strains*). We estimated the magnitude of this effect by fitting the log-affinity-fluorescence relationship for these variants, including an effect of the RE, using the equation

$$F = L + \frac{U - L}{1 + e^{-\frac{\log(K_A) + d}{a}}} \quad (8)$$

where $d$ is the difference in $log(K_A)$ for GA-bound variants and $a$ determines the steepness of the curve. The best-fit estimate is that the FS mutations reduce the $K_A$ of GA across DBD variants by 0.95 ± SE 0.12 $\log_{10}(\mu M^{-2})$. The genetic score of a complex in the RFA model is linearly related to its affinity, so we used this estimate to adjust the genetic score of all variants on GA and used the fluorescence predicted by the model after this adjustment (Extended Data Fig. 3d). The resulting transformation increased the number of inferred active GA RE variants from 5 to 75 in the AncSR1 background and from 449 to 1,172 in the AncSR2 background. It also increased the fluorescence of the wild type AncSR2 protein on GA to $F = -4.28$, which is closer to the measured fluorescence of AncSR2 wild type on SRE, as predicted by [10,11].

*Accuracy of predicted functional classification*

To classify protein-RE variants with predicted fluorescence as functional or non-functional, we performed a nonparametric bootstrap test to address model prediction error: we concatenated residuals across the 10 RFA cross-validation models, then sampled residuals from within an interval of ±0.1 fluorescence units from the inferred fluorescence of each complex (Extended Data Fig. 3e). *p*-values were calculated as the proportion of bootstrap samples ($n = 250$) with fluorescence greater than or equal to that of the wild type complex.

To assess the accuracy of this functional classification, we analyzed the prediction error from the CV fits with the same regularization strength as the final models, which indicate how well the full model predicts unseen data. The AncSR2 CV models predicted the held-out data reasonably well, with mean $R^2 = 0.79$ for all variants and $R^2 = 0.75$ for active variants across the 10 CV fits (Extended Data Fig. 3e, right). However, prediction was considerably worse for the AncSR1 CV models, with mean $R^2 = 0.20$ for all variants and $R^2 = 0.07$ for active variants (Extended Data Fig. 3e, left). This can in large part be attributed to a bias towards underprediction observed in the AncSR1 CV models, which likely results from the fact that the vast majority of variants in this protein background are at the lower bound of fluorescence; genetic states that may be beneficial for binding in some genetic contexts are thus only seen at the lower bound, resulting in a negative bias in their inferred effects (Extended Data Fig. 3e). The AncSR2 CV models also exhibit a slight underprediction bias, but the effect is much less severe because there are many more active variants in this protein background. The prediction error and bias from the models are taken into account in later analyses.

*Protein-RE genotype networks*

To describe the effects of coevolution, we built joint protein-DNA genotype networks. We considered two functional protein-RE complexes as mutational neighbors if they differed by a single amino acid in the protein *or* a single nucleotide in the RE. For example, the genotypes EGKA:GT and EGKV:GT are neighbors by mutations in the protein, and the genotypes AAAI:AA and AAAI:TA are neighbors due to mutations in the RE. In this network, promiscuous genotypes (such as AAAI) are represented as two separate nodes, one for each complex. Network analyses and visualization were done as in the protein genotype networks.

Although both the protein and the RE can substitute in this model, we calculated the length of evolutionary trajectories by including only changes in the protein's amino acid sequence, in order to fairly compare the length of these trajectories to those in the protein-only network. We therefore adjusted the total number of protein+DNA substitutions in any trajectory using the average fraction of all steps on the network that involve changes in the protein sequence. To estimate this fraction, we first computed for every protein-RE complex *g* the fraction of one-mutation neighbors that change the protein sequence ($f_{AA,g}$). We ran a Markov chain starting from the set of all genotypes in the network, and at each step ($k$) we computed the fraction of all possible steps in the network that are amino acid changes ($N_{AA,k}$) as:

$$N_{AA,k} = \sum_{g} f_{AA,g} \times \pi_{(k)g} \tag{19}$$

where $g$ iterates over all complexes and $\pi_{(k)}$ is the vector of probabilities of genotypes after $k$ steps of the Markov chain. We took the average fraction across trajectory lengths and used this to scale the length of any evolutionary trajectory in the coevolution network as the expected number of amino acid steps on that trajectory.

*Protocol: Combinatorial library assembly and bacterial transformation*

This protocol aims to obtain ~100X coverage per barcoded DBD library (*i.e.* ~$2\times10^7$ cfu). The protocol was repeated for each of the 32 barcoded libraries.

1. Perform second strand synthesis on 5 pmol of library oligo using Q5 DNA polymerase in a 10-µL PCR reaction. Denature at 95°C for 80 s, extend at 50°C for 30 s, anneal at 72°C for 3 min. 30 s.
2. Assemble the library oligo into the plasmid backbone using the BsaI-HFv2 Golden Gate Assembly Kit (NEB). For one 50-µL reaction, use 0.4 µL of double stranded library oligo (0.2 pmol) from the previous step and 0.1 pmol of pDBD2.1 AncSR1 or AncSR2 vector with BsaI cut sites. Perform 2 50-µL reactions per library. Incubate reactions at 37°C for 1 hr, then 50°C for 5 min. to inactivate the enzyme.
3. Pool the 50-µL reactions and purify using the Zymo Clean & Concentrator-5 kit. Elute in 8 µL dd$H_2O$ and chill on ice.
4. Transform 4 µL of assembled pDBD2.1 library into Invitrogen ElectroMAX DH5α-E competent cells following the manufacturer's protocol. Electroporate at 1.7 kV in 1-mm cuvettes. Immediately add 1 mL of preheated SOC to the cuvette and transfer the mixture to a culture tube, using a P1000 tip inserted into a P20 tip.
5. Recover the cells for 1 hour at 37°C, 225 rpm.
6. Take 10 µL of the recovered culture and dilute with 1 mL LB. Take 10 µL of this culture and dilute again with 1 mL LB. Plate 100 µL of this culture on LB + 100 µg/mL carbenicillin. This constitutes a $2\times10^5$-fold dilution.
7. Transfer the rest of the recovered cells to 25 mL LB + 100 µg/mL carbenicillin in a culture flask. Grow overnight at 37°C, 225 rpm.
8. The following morning, take 750 µL of the overnight culture to make 20% glycerol stocks. Maxiprep the remaining culture using the GenElute HP Plasmid DNA Maxiprep Kit. Elute DNA in 3 mL dd$H_2O$ and concentrate DNA to 1.2 mL by ethanol precipitation. Measure concentration with NanDrop (should be ~500 ng/µL).
9. Count colonies on the serial dilution plate to estimate the number of colony forming units (cfu) obtained from transformation. Pick several colonies for Sanger sequencing to verify that library assembly was successful.

*Protocol: Yeast transformation*

This protocol is for transforming one barcoded DBD library into a yeast RE reporter strain. We were able to transform three libraries at a time with two people working together; this is reflected in the media and buffer volumes. The protocol aims to obtain ~50X coverage per library (*i.e.* ~$1\times10^7$ cfu).

*Media and Buffers:*

2L 2X YPD

- 20 g Yeast extract
- 40 g Peptone
- Add dd$H_2O$ to 1.9 L in a 2 L flask and autoclave
- Add 100 mL filter-sterilized 40% dextrose

600 mL Buffer E (1M sorbitol, 1mM $CaCl_2$)

- 109.2 g sorbitol
- 0.087 g $CaCl_2$ [87 mg] (or 0.113 g $CaCl_2.2H_2O$ [113 mg])
- Bring to volume with dd$H_2O$
- Filter sterilize
- Store at 4°C

250 mL Buffer C (10X TE, 1M LiAc, 1M DTT), prepared fresh

- 2.5 mL 1 M Tris-Cl pH 8.0
- 500 uL 0.5 M EDTA pH 8.0
- 25 mL 1 M LiOAc
- 2.5 mL 1 M DTT
- 219.5 mL dd$H_2O$
- Filter sterilize into 50-mL conical tubes

600 mL recovery media (1M sorbitol in YPD)

- 109.26 g sorbitol
- Bring to 600mL with YPD, filter sterilize.

600 mL sterile water, chilled to 0°C

<u>Day 1</u>

1. Streak yeast glycerol stocks on a YPD plate.

<u>Day 2 (at least 15h before day 3)</u>

2. Inocubate 150 mL YPD with a single colony of each RE yeast strain in 1 L flask; incubate at 30°C, 225 rpm overnight.

Day 3

*Preparation*

Chill centrifuge to 4°C

3. Read overnight $OD_{600}$.
4. Aliquot 400 $OD_{600}$-mL in 50-mL conical tubes; spin 3 kg 2 min; discard supernatant.
5. Resuspend in 800 mL YPD in two 2.5 L flasks.
6. Incubate at 30°C, 225 rpm until $OD_{600}$ > 4 (~4h)
7. Chill autoclaved water, Buffer E, 2-mm cuvettes, and prepare 250 mL fresh Buffer C.
8. Spin down cells in 8 50-mL conical tubes (two spins per tube); spin 3 kg 2 min; discard supernatant.
9. Resuspend and pool cells in 90 mL ice-cold dd$H_2O$ in two 50-mL conicals. Spin 3 kg 2 min. Discard supernatant.
10. Wash once more with 45 mL ice-cold dd$H_2O$ each tube.
11. Resuspend each in 45 mL ice-cold Buffer E; spin 3 kg 2 min; discard supernatant.
12. Resuspend both tubes in 160 mL Buffer C and add them to a 1 L culture flask. Incubate for 30 min at 30°C, 225 rpm.
13. Pellet the cells in two 50-mL conicals; decant supernatant. Wash once with ice-cold Buffer E (45 mL per tube); spin 3 kg 2 min. Decant supernatant.
14. Resuspend pellets in remaining volume of Buffer E; pool and add Buffer E for a final volume of 4.8 mL cells + DNA per library (add ~1.5 mL Buffer E per strain to resuspend). Resuspend using a 25 mL pipette and minimize cell loss on sides of pipette/tubes.

*Electroporation*

15. Mix 600 uL of plasmid library to tube with cells (at least 288 µg of DNA).
16. Add 171.2 mL of recovery media in an empty 1-L flask.
17. Aliquot 400 µL of cell/plasmid mixture in each chilled 2-mm electroporation cuvette (~12 cuvettes per library).
18. Electroporate at 2.5kV; immediately add 1 mL of recovery media at RT.
    - Add the recovery media gently and pipette up and down to bring the cells on the bottom of the cuvette to the surface.
19. Pool reactions together in a 1-L flask with an additional 171.2 mL of recovery media. Rinse each cuvette with an additional 1 mL of recovery media (final volume: 200 mL).
20. Incubate the transformed cells at 30°C, 225 rpm for 2 hr.
21. Make $10^{-4}$ and $10^{-5}$ serial dilution plates (YPD + 200 µg/mL G418) to estimate transformation yield.

*Passage and storage*

22. Aliquot recovery into 50-mL conicals.

23. Spin 3kg 2min room temperature; discard supernatant.
24. Resuspend to 400 mL YPD + 200 µg/mL G418 + 25 µg/mL chlor in a 2.5-L flask.
25. Incubate at 30°C 225 rpm until saturation, approximately 24 hr.

Day 4

26. Make 20 1-mL 25% glycerol stocks with 200 $OD_{600}$-mL of yeast; flash freeze in liquid $N_2$; store at -80°C.

**Supplementary Table 1: Synonymous RE barcodes (REBCs)**

Sequences of synonymous mutations used to barcode the DBD libraries to associate them with RE strains. NNS, variable sites in the DMS experiment.

| RE barcode ID | RE barcode sequence | RE |
|---|---|---|
| AncSR1 REBC 1 | GGTGTATGGTCATGTNNSNNSTGTNNSNNS | SRE |
| AncSR1 REBC 2 | GGGGTTTGGTCGTGTNNSNNSTGTNNSNNS | GA |
| AncSR1 REBC 3 | GGGGTTTGGAGTTGTNNSNNSTGTNNSNNS | ERE |
| AncSR1 REBC 4 | GGTGTATGGAGCTGTNNSNNSTGTNNSNNS | AC |
| AncSR1 REBC 5 | GGCGTTTGGTCATGCNNSNNSTGTNNSNNS | AG |
| AncSR1 REBC 6 | GGAGTATGGTCGTGCNNSNNSTGTNNSNNS | AT |
| AncSR1 REBC 7 | GGTGTCTGGAGTTGCNNSNNSTGTNNSNNS | CA |
| AncSR1 REBC 8 | GGCGTGTGGAGCTGCNNSNNSTGTNNSNNS | CC |
| AncSR1 REBC 9 | GGCGTCTGGTCATGTNNSNNSTGCNNSNNS | CG |
| AncSR1 REBC 10 | GGAGTGTGGTCGTGTNNSNNSTGCNNSNNS | CT |
| AncSR1 REBC 11 | GGCGTGTGGAGTTGTNNSNNSTGCNNSNNS | GC |
| AncSR1 REBC 12 | GGAGTCTGGAGCTGTNNSNNSTGCNNSNNS | GG |
| AncSR1 REBC 13 | GGTGTGTGGTCATGCNNSNNSTGCNNSNNS | TA |
| AncSR1 REBC 14 | GGGGTCTGGTCGTGCNNSNNSTGCNNSNNS | TC |
| AncSR1 REBC 15 | GGAGTTTGGAGTTGCNNSNNSTGCNNSNNS | TG |
| AncSR1 REBC 16 | GGGGTATGGAGCTGCNNSNNSTGCNNSNNS | TT |
| AncSR2 REBC 1 | GTGTTGACGTGCNNSNNSTGCNNSNNS | SRE |
| AncSR2 REBC 2 | GTTCTTACGTGCNNSNNSTGCNNSNNS | GA |
| AncSR2 REBC 3 | GTATTAACATGCNNSNNSTGCNNSNNS | ERE |
| AncSR2 REBC 4 | GTCCTCACATGCNNSNNSTGCNNSNNS | AC |
| AncSR2 REBC 5 | GTACTTACATGTNNSNNSTGCNNSNNS | AG |
| AncSR2 REBC 6 | GTGCTCACCTGTNNSNNSTGCNNSNNS | AT |
| AncSR2 REBC 7 | GTTCTGACTTGTNNSNNSTGCNNSNNS | CA |
| AncSR2 REBC 8 | GTGCTTACATGCNNSNNSTGTNNSNNS | CC |
| AncSR2 REBC 9 | GTCTTAACCTGCNNSNNSTGTNNSNNS | CG |
| AncSR2 REBC 10 | GTTCTCACCTGCNNSNNSTGTNNSNNS | CT |
| AncSR2 REBC 11 | GTCCTGACTTGCNNSNNSTGTNNSNNS | GC |
| AncSR2 REBC 12 | GTATTGACGTGTNNSNNSTGTNNSNNS | GG |
| AncSR2 REBC 13 | GTTCTAACGTGTNNSNNSTGTNNSNNS | TA |
| AncSR2 REBC 14 | GTCCTTACCTGTNNSNNSTGTNNSNNS | TC |
| AncSR2 REBC 15 | GTGTTAACTTGTNNSNNSTGTNNSNNS | TG |
| AncSR2 REBC 16 | GTACTCACTTGTNNSNNSTGTNNSNNS | TT |

**Supplementary Table 2: Library transformation and enrichment sort statistics**

Library transformation yields, glycerol stock recovery rates, and number of cells sorted for enrichment sorts.

| DBD background | RE strain | RE barcode | Bacterial transfor-mation cfu (x10^7) | Yeast transfor-mation cfu (x10^7) | Top-up yeast transfor-mation cfu (x10^7) | Total yeast transfor-mation cfu (x10^7) | Enrich-ment sort batch | Glycerol stock recovery cfu (x10^7) | Enrich-ment sort GFP– cell count | Enrich-ment sort GFP+ cell count | Enrich-ment sort GFP+ proportion |
|---|---|---|---|---|---|---|---|---|---|---|---|
| AncSR1 | SRE | 1 | 5.67 | 3.09 | | 3.09 | 1 | 19.4 | 24375871 | 625635 | 0.025 |
| AncSR1 | GA | 2 | 2.01 | 2.82 | | 2.82 | 1 | 29.6 | 24800681 | 435760 | 0.017 |
| AncSR1 | ERE | 3 | 2.38 | 2.58 | | 2.58 | 1 | 25.4 | 24681039 | 405514 | 0.016 |
| AncSR1 | AC | 4 | 2.64 | 1.73 | | 1.73 | 1 | 10.6 | 24592905 | 528709 | 0.021 |
| AncSR1 | AG | 5 | 2.07 | 3.25 | | 3.25 | 1 | 25.8 | 24662980 | 408313 | 0.016 |
| AncSR1 | AT | 6 | 3.39 | 3.8 | | 3.8 | 1 | 25.8 | 24806774 | 472083 | 0.019 |
| AncSR1 | CA | 7 | 1.56 | 2.07 | | 2.07 | 1 | 14.4 | 24352862 | 726379 | 0.029 |
| AncSR1 | CC | 8 | 1.08 | 2.9 | | 2.9 | 1 | 27.8 | 24658946 | 448824 | 0.018 |
| AncSR1 | CG | 9 | 1.66 | 0.85 | 1.1 | 1.95 | 5/6 | 18.4 | 24326839 | 858760 | 0.034 |
| AncSR1 | CT | 10 | 2.37 | 2.34 | | 2.34 | 5/6 | 25.6 | 24736293 | 379479 | 0.015 |
| AncSR1 | GC | 11 | 1.97 | 2.26 | | 2.26 | 5/6 | 26.4 | 24336612 | 744771 | 0.03 |
| AncSR1 | GG | 12 | 1.38 | 1.32 | | 1.32 | 5/6 | 33.6 | 24841745 | 285197 | 0.011 |
| AncSR1 | TA | 13 | 1.86 | 0.69 | 1.29 | 1.98 | 5/6 | 10.4 | 24748789 | 329854 | 0.013 |
| AncSR1 | TC | 14 | 1.1 | 2.16 | | 2.16 | 5/6 | 20.6 | 24558731 | 587774 | 0.023 |
| AncSR1 | TG | 15 | 1.22 | 0.91 | 1.01 | 1.92 | 2/6 | 15.6 | 24945619 | 348916 | 0.014 |
| AncSR1 | TT | 16 | 3.07 | 2.54 | | 2.54 | 2/6 | 16.4 | 25226917 | 437954 | 0.017 |
| AncSR2 | SRE | 1 | 5.24 | 1.79 | | 1.79 | 4 | 17.8 | 24220370 | 847039 | 0.034 |

| | | | | | | | | | | | |
|---|---|---|---|---|---|---|---|---|---|---|---|
| AncSR2 | GA | 2 | 3.79 | 1.37 | | 1.37 | 4 | 25 | 24825719 | 391276 | 0.016 |
| AncSR2 | ERE | 3 | 5.1 | 1.5 | | 1.5 | 4 | 17.4 | 24694083 | 425418 | 0.017 |
| AncSR2 | AC | 4 | 3.54 | 0.7 | 0.87 | 1.57 | 4 | 9.8 | 24602087 | 533981 | 0.021 |
| AncSR2 | AG | 5 | 4.6 | 0.43 | 0.9 | 1.33 | 4 | 8.8 | 24690832 | 434945 | 0.017 |
| AncSR2 | AT | 6 | 4.3 | 1.6 | | 1.6 | 4 | 1.6 | 24686331 | 421661 | 0.017 |
| AncSR2 | CA | 7 | 2.2 | 1.17 | | 1.17 | 4 | 1.6 | 24407887 | 777213 | 0.031 |
| AncSR2 | CC | 8 | 2.8 | 1.35 | | 1.35 | 4 | 2.8 | 24637616 | 409661 | 0.016 |
| AncSR2 | CG | 9 | 6.9 | 1.98 | | 1.98 | 3 | 18.6 | 24399903 | 903819 | 0.036 |
| AncSR2 | CT | 10 | 4.5 | 1.16 | | 1.16 | 3 | 4.6 | 24518676 | 796209 | 0.031 |
| AncSR2 | GC | 11 | 5.4 | 0.97 | | 0.97 | 3 | 21 | 24214644 | 940935 | 0.037 |
| AncSR2 | GG | 12 | 5.1 | 2.52 | | 2.52 | 3 | 18 | 24511230 | 679816 | 0.027 |
| AncSR2 | TA | 13 | 1.46 | 1.08 | | 1.08 | 3 | 8.2 | 24606769 | 673875 | 0.027 |
| AncSR2 | TC | 14 | 12.9 | 1.3 | | 1.3 | 3 | 18 | 24040623 | 1105703 | 0.044 |
| AncSR2 | TG | 15 | 4.6 | 1.5 | | 1.5 | 3 | 10.8 | 24441448 | 617178 | 0.025 |
| AncSR2 | TT | 16 | 2.7 | 2.03 | | 2.03 | 3 | 20.8 | 24619287 | 854539 | 0.034 |

**Supplementary Table 3: Binned sort statistics**

Glycerol stock recovery rates and number of cells sorted for binned sorts.

| Binned sort replicate | Background | Libraries | Enrich-ment sort batch | Glycerol stock recovery cfu (x10^7) | Glycerol stock recovery cfu per GFP+ cells sorted | Cells sorted, bin 1 | Cells sorted, bin 2 | Cells sorted, bin 3 | Cells sorted, bin 4 | Cells sorted, total |
|---|---|---|---|---|---|---|---|---|---|---|
| 1 | AncSR1 | 1-8 | 1 | 2.8 | 6.9 | 68112878 | 86824472 | 5447744 | 3638776 | 164023870 |
| 1 | AncSR1 | 15-16 / 9-14 | 2 / 5 | 2.3 / 1.2 | 29.2 / 3.8 | | | | | |
| 1 | AncSR2 | 1-8 | 4 | 10.2 | 24 | | | | | |
| 1 | AncSR2 | 9-16 | 3 | 4 | 6.1 | | | | | |
| 2 | AncSR1 | 1-8 | 1 | 1.8 | 4.4 | 71873842 | 75663838 | 5380470 | 3116191 | 156034341 |
| 2 | AncSR1 | 9-16 | 6 | 29.4 | 90.3 | | | | | |
| 2 | AncSR2 | 1-8 | 4 | 15.6 | 36.8 | | | | | |
| 2 | AncSR2 | 9-16 | 3 | 5.8 | 8.8 | | | | | |
| 3 | AncSR1 | 1-8 | 1 | 1.8 | 4.4 | 66662146 | 87779826 | 7047242 | 4303413 | 165792627 |
| 3 | AncSR1 | 9-16 | 6 | 4.8 | 14.7 | | | | | |
| 3 | AncSR2 | 1-8 | 4 | 19 | 44.8 | | | | | |
| 3 | AncSR2 | 9-16 | 3 | 7.2 | 11 | | | | | |
| 4 | AncSR1 | 9-16 | 6 | 4.6 | 14.1 | 16233786 | 12616013 | 822095 | 579424 | 30251318 |

**Supplementary Table 4: Binned sort sequencing statistics**

Estimated number of reads per cell across libraries, bins, and replicates. Libraries for AncSR1 REBC 9–14 and REBC 15–16 had very low and high sequencing depths, respectively, in replicate 1, so we repeated the enrichment sort for these libraries and used them for bins sort replicates 2–4.

| Background | REBC | Rep 1 reads/cell | Rep 2 reads/cell | Rep 3 reads/cell | Rep 4 reads/cell |
|---|---|---|---|---|---|
| AncSR1 | 1 | 0.86 | 0.56 | 1.35 | |
| AncSR1 | 2 | 1.16 | 0.8 | 1.83 | |
| AncSR1 | 3 | 0.91 | 0.62 | 1.56 | |
| AncSR1 | 4 | 1.9 | 1.29 | 3.27 | |
| AncSR1 | 5 | 2.37 | 1.57 | 3.87 | |
| AncSR1 | 6 | 2.39 | 1.68 | 3.81 | |
| AncSR1 | 7 | 1.41 | 0.93 | 2.4 | |
| AncSR1 | 8 | 2.11 | 1.46 | 3.57 | |
| AncSR1 | 9 | 0.06 | 0.33 | 1.2 | 1.55 |
| AncSR1 | 10 | 0.04 | 1.26 | 4.04 | 1.4 |
| AncSR1 | 11 | 0.06 | 0.3 | 1.01 | 1.89 |
| AncSR1 | 12 | 0.05 | 1.17 | 3.78 | 2.01 |
| AncSR1 | 13 | 0.04 | 0.64 | 1.99 | 1.18 |
| AncSR1 | 14 | 0 | 0.36 | 1.18 | 0.98 |
| AncSR1 | 15 | 8.97 | 1.09 | 3.59 | 2 |
| AncSR1 | 16 | 4.7 | 0.58 | 1.86 | 1.42 |
| AncSR2 | 1 | 1.38 | 0.86 | 4.43 | |
| AncSR2 | 2 | 0.84 | 0.57 | 2.74 | |
| AncSR2 | 3 | 1.41 | 0.88 | 4.77 | |
| AncSR2 | 4 | 1.07 | 0.67 | 3.7 | |
| AncSR2 | 5 | 1.01 | 0.63 | 3.39 | |
| AncSR2 | 6 | 0.99 | 0.64 | 3.08 | |
| AncSR2 | 7 | 1.39 | 0.83 | 4.63 | |
| AncSR2 | 8 | 1.64 | 1.01 | 5.65 | |
| AncSR2 | 9 | 4.26 | 2.15 | 12.19 | |
| AncSR2 | 10 | 0.77 | 0.44 | 2.16 | |
| AncSR2 | 11 | 1.55 | 0.84 | 4.38 | |
| AncSR2 | 12 | 0.98 | 0.57 | 2.67 | |
| AncSR2 | 13 | 0.28 | 0.16 | 0.77 | |
| AncSR2 | 14 | 0.58 | 0.32 | 1.58 | |
| AncSR2 | 15 | 0.61 | 0.35 | 1.7 | |
| AncSR2 | 16 | 0.41 | 0.23 | 1.11 | |
